# Gradual evolution of allopolyploidy in *Arabidopsis suecica*

**DOI:** 10.1101/2020.08.24.264432

**Authors:** Robin Burns, Terezie Mandáková, Joanna Gunis, Luz Mayela Soto-Jiménez, Chang Liu, Martin A. Lysak, Polina Yu. Novikova, Magnus Nordborg

**Affiliations:** Gregor Mendel Institute, Austrian Academy of Sciences, Vienna BioCenter, Vienna, Austria; CEITEC - Central European Institute of Technology, and Faculty of Science, Masaryk University, Brno, Czech Republic; Institute of Biology, University of Hohenheim, Garbenstrasse 30, 70599 Stuttgart, Germany; VIB-UGent Center for Plant Systems Biology, Ghent, Belgium; Department of Chromosome Biology, Max Planck Institute for Plant Breeding Research, Cologne, Germany

## Abstract

The majority of diploid organisms have polyploid ancestors. The evolutionary process of polyploidization (and subsequent re-diploidization) is poorly understood, but has frequently been conjectured to involve some form of “genome shock” — partly inspired by studies in crops, where polyploidy has been linked to major genomic changes such as genome reorganization and subgenome expression dominance. It is unclear, however, whether such dramatic changes would be characteristic of natural polyploidization, or whether they are a product of domestication. Here, we study polyploidization in *Arabidopsis suecica* (n = 13), a post-glacial allopolyploid species formed via hybridization of *A. thaliana* (n = 5) and *A. arenosa* (n = 8). We generated a chromosome-level genome assembly of *A. suecica* and complemented it with polymorphism and transcriptome data from multiple individuals of all species. Despite a divergence of ∼6 Mya between the two ancestral species and appreciable differences in their genome composition, we see no evidence of a genome shock: the *A. suecica* genome is highly colinear with the ancestral genomes, there is no subgenome dominance in expression, and transposable element dynamics appear to be stable. We do, however, find strong evidence for changes suggesting gradual adaptation to polyploidy. In particular, the *A. thaliana* subgenome shows upregulation of meiosis-related genes, possibly in order to prevent aneuploidy and undesirable homeologous exchanges that are frequently observed in experimentally generated *A. suecica*, and the *A. arenosa* subgenome shows upregulation of cyto-nuclear related processes, possibly in response to the new cytoplasmic environment of *A. suecica,* with plastids maternally inherited from *A. thaliana*.

## Introduction

Ancient polyploidization or whole-genome duplication is a hallmark of most higher-organism genomes^1, 2^, including our own^3, 4^. While most of these organisms are now diploid and show only traces of polyploidy, there are many examples of recent polyploidization, in particular among flowering plants^5–9^. These examples are important because they allow us to study the process of polyploidization, rather than just inferring that it happened and trying to understand its evolutionary importance.

Wide-spread naturally occuring polyploid hybrids (i.e. allopolyploids), such as *Capsella bursa-pastoris* (Shepherd’s Purse)^10–12^, *Trifolium repens* (white clover)^13^, *Brachypodium hybridum*^14, 15^*, Arabidopsis kamchatica*^16^, *Mimulus peregrinus*^17^*, Tragopogon miscellus* and *T. mirus*^18^, demonstrate that natural polyploid species can quickly become successful, and even be deemed invasive^19^. Regardless of their eventual evolutionary success, new allopolyploid species face numerous challenges, ranging from those on a population level, such as bottlenecks^13, 20^ and competition with their diploid progenitors^21^, to those on a genomic level, such as chromosome segregation^22–24^ and changes to hybrid genome structure (e.g. chromosomal structural variants and aneuploidy^25^) and genome regulation (e.g. subgenome expression dominance^26^ and the regulation of transposable elements^27^) — phenomena which may be enhanced by genomic conflicts between the newly merged subgenomes, leading to a “genome shock”^28^. In agreement with this, genomic and transcriptomic changes tied to the hybridization of two (or more) diverged genomes have been reported in resynthesized polyploids of wheat^29–35^, *Brassica napus*^36–38^ and cotton^39, 40–37, 41, 42^ (although resynthesized cotton appears genetically stable^43^).

The long-term importance of such rapid changes is less clear. For example, the transposable element transcription and mobilization observed in resynthesized wheat^33, 44–46^, is not reflected in the genome sequence of cultivated wheat^47^. However, other cultivated crop genomes, for example cotton, show instances of large structural rearrangements^5, 48–50^, biased gene loss^51^, a spreading and proliferation of centromere repeats between subgenomes^52^ and changes to the 3D genome structure^53^. Strawberry^6^, peanut^8^ and the mesopolyploids *B. rapa*^54^ and maize^55^ show evidence of subgenome dominance, while wheat^56^, cotton^51^ and *B. napus*^57^ do not. The reasons for these differences are not understood.

An even greater source of uncertainty is whether allopolyploid crops are representative of natural polyploidization. Domestication is frequently associated with very strong “artifical” selection, which can dramatically alter the fitness landscape^58–62^. For example, large structural variants have been linked to favourable agronomic traits^63–65^. In addition, polyploid crops are generally quite recent, evolutionarily speaking.

Turning to non-domesticated species, genomic changes have been reported in natural allopolyploids like the ∼80 years old *Tragopogon miscellus*^66, 67^, the ∼140 years old *Mimulus pergrinus*^17^, and *Spartina anglica*^68^, which likely originated at the end of the 19th century — however, these examples are extremely recent and are more in line with the reported genomic changes in the resynthesized allopolyploids. Older natural allopolyploids, on the other hand, generally do not show signs of genomic changes after allopolyploidy. Examples of these include: white clover^13^, *C. bursa-pastoris*^12, 69^, *A. kamchatica*^16, 70^*, B. hybridium*^14^ and the gymnosperm *Ephedra*^71^.

Here we focus on an allopolyploid comparable in age to these examples, the highly selfing^72^, *A. suecica* (2n = 4x = 26), formed through the hybridization of *A. thaliana* (2n = 10) and *A. arenosa* (2n = 2x/4x = 16/32), circa 16 kya, during the Last Glacial Maximum^20^ and now widely established in northern Fennoscandia (Fig. 1a). The ancestral species diverged around 6 Mya^73^, and, based on mitochondrial and chloroplast sequences, it is clear that *A. thaliana* is the maternal and *A. arenosa* the paternal parent of the hybrid^74^, a scenario also supported by the fact that *A. arenosa* itself is a ploidy-variable species, so that *A. suecica* could readily be generated through the fertilization of an unreduced egg cell (2n = 2x) from *A. thaliana* by a sperm cell (n = 2x) from autotetraploid *A. arenosa*^20, 75^. We have previously shown that, although *A. suecica* shows clear evidence of a genetic bottleneck^20^, it shares most of its variation with the ancestral species, demonstrating that the species was formed through a hybridization and polyploidization process that involved many crosses and individuals. In order to study genomic change in *A. suecica*, we used long-read sequencing to generate a high-quality, chromosome-level assembly of a single individual, taking advantage of the fact that *A. suecica*, like *A. thaliana*, is highly selfing, making it possible to sequence naturally inbred individuals. The genome sequence was complemented by a partial assembly of a tetraploid outcrosser *A. arenosa*, and by short-read genome and transcriptome sequencing data from many individuals of all three species — including “synthetic” *A. suecica* generated *de novo* in laboratory crosses.

**Figure 1.**
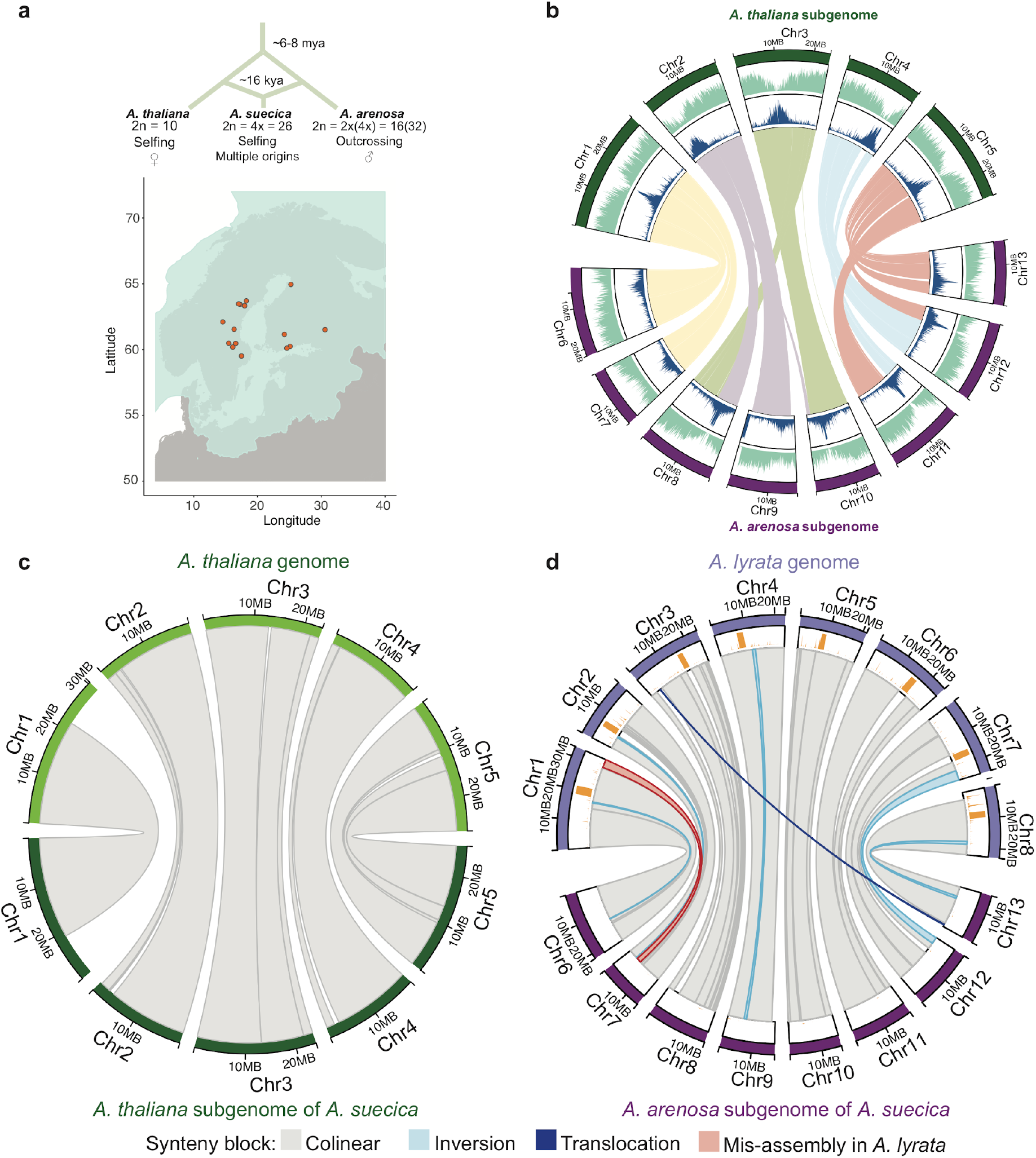
The genome of *A. suecica* is largely colinear with the ancestral genomes. **a** Schematic depicting the origin of *A. suecica* and its current distribution in the relation to the ice cover at the last glacial maximum (LGM). **b** The chromosome-level assembly of the *A. suecica* genome with inner links depicting syntenic blocks between the *A. thaliana* and *A. arenosa* subgenomes of *A. suecica*. The blue histogram represents the distribution of TEs along the genome and the green histogram corresponds to the distribution of protein-coding genes. **c** Synteny of the *A. thaliana* subgenome of *A. suecica* to the *A. thaliana* TAIR10 reference. In total 13 colinear synteny blocks were found. **d** Synteny of the *A. arenosa* subgenome to *A. lyrata*. In total 40 synteny blocks were found, 33 of which were colinear. Of the remaining 7 blocks, 5 represent inversions in the *A. arenosa* subgenome of *A. suecica* compared to *A. lyrata*, 1 is a translocation, and 1 corresponds to a previously reported mis-assembly in the *A. lyrata* genome^77^. Orange bars represent a density plot of missing regions (“N” bases) in the *A. lyrata* genome.

## Results and discussion

### 1. The genome is conserved

We assembled a reference genome from a naturally inbred (i.e. the species is self-compatible^20, 72^) *A. suecica* accession (“ASS3”), using 50x long-read PacBio sequencing (PacBio RS II). The absence of heterozygosity and the substantial (∼11.6%) divergence between the subgenomes greatly facilitated the assembly. In contrast, assembling even a diploid genome of the outcrosser *A. arenosa* is complicated by high heterozygosity (nucleotide diversity around 3.5%^76^) coupled with a relatively high level of repetitive sequences (compared to the gene-rich *A. thaliana* genome). Our attempt to assemble a tetraploid *A. arenosa* individual, the result of which is also included here in addition to the genome of *A. suecica*, led to a very fragmented assembly of 3,629 contigs with an N50 of 331 Kb. In contrast, the *A. suecica* assembly has an N50 contig size of 9.02 Mb. The assembled contigs totaled 276 Mb (∼90% of the 305 Mb genome size estimated by flow cytometry — see Supplementary Fig. 1; ∼88% of the 312Mb genome size estimated by kmer analysis). Contigs were placed into scaffolds using high-coverage chromosome conformation capture (HiC) data and by using the reference genomes of *A. thaliana* and *A. lyrata* (here the closest substitute for *A. arenosa*) as guides. This resulted in 13 chromosome-scale scaffolds (Supplementary Fig. 2a). The placement and orientation of each contig within a scaffold was confirmed and corrected using a genetic map for *A. suecica* (see Methods, Supplementary Fig. 3, Supplementary Fig. 4). The resulting chromosome-level assembly (Fig. 1b) contains 262 Mb, and has an N50 scaffold size of 19.59 Mb. The five chromosomes of the *A. thaliana* subgenome and the eight chromosomes of the *A. arenosa* subgenome sum to 119 Mb and 143 Mb, respectively.

Approximately 108 and 135 Mb of the *A. thaliana* and *A. arenosa* subgenomes of *A. suecica* are in large blocks syntenic to the genomes of the ancestral species: 13 and 40 blocks, respectively (Fig. 1c,d). The vast majority of these syntenic blocks are themselves also colinear, with the exception of five small-scale inversions (∼4.5 Mb) and one translocation (∼244 Kb) on the *A. arenosa* subgenome— which may well (indeed probably do) reflect differences between *A. lyrata* and *A. arenosa*, two highly polymorphic species separated by about a million years^73, 76^. We also corrected for the described^77^ mis-assembly in the *A. lyrata* reference genome using our genetic map. Overall we find that approximately 93% of the *A. suecica* genome is syntenic to the ancestral genomes, the 13 chromosomes of *A. suecica* having remained almost completely colinear (Fig. 1c,d). This highlights the conservation of the *A. suecica* genome and contrasts with the major rearrangements that have been observed in several resynthesized polyploids^29, 32, 34, 36^ and some crops^48, 50, 78^. Interestingly, major rearrangements have also been observed in synthetic *A. suecica*^79^, and we see clear evidence of aneuploidy in ours — a topic to which we shall return.

A total of 45,585 protein-coding genes were annotated for the *A. suecica* reference, of which 22,232 and 23,353 are located on the *A. thaliana* and *A. arenosa* subgenomes, respectively. We assessed completeness of the genome assembly and annotation with the BUSCO set for eudicots and found 2088 (98.4%) complete genes for both the *A. thaliana* and *A. arenosa* subgenomes (Supplementary Fig. 5c,d). Of the protein-coding genes, 18,023 had a one-to-one orthology between the subgenomes of *A. suecica* and 16,999 genes were conserved in a 1:1:1:1 relationship for both subgenomes of *A. suecica* and the ancestral species (using *A. lyrata* as a substitute for *A. arenosa*) (Supplementary Data 2, Supplementary Fig. 5b). We functionally annotated lineage-specific genes in *A. suecica* (i.e. genes in *A. suecica* without a reciprocal best blast hit to *A. thaliana* or *A. lyrata*) using InterPro, and only found significant enrichment in *A. thaliana* subgenome of *A. suecica* for two GO terms (GO:0008234 and GO:0015074), both of which are associated with repeat content (Supplementary Data 2). Ancestral genes not found in the *A. suecica* genome annotation were overrepresented for functional categories of plant defense response. However, checking coverage for these genes by mapping the raw *A. suecica* whole-genome resequencing data to the ancestral genomes did not confirm their loss, suggesting rather misassembly or misannotation, which is expected due to the repetitive and highly polymorphic nature of R-genes in plants.

### 2. The rDNA clusters are highly variable

In eukaryotic genomes, genes encoding ribosomal RNA (rRNA) occur as tandem arrays in rDNA clusters. The 45S rDNA clusters are particularly large, containing hundreds or thousands of copies, spanning millions of base pairs^80^. The nucleolus, the site of pre-ribosome assembly, forms at these clusters, but only if they are actively transcribed, and it was observed long ago that only one parent’s rDNA tended to be involved in nucleolus formation in inter-specific hybrids, a phenomenon known as “nucleolar dominance”^81–84^. In *A. suecica,* it was observed that the rDNA clusters inherited from *A. thaliana* were silenced^85–87^, and structural changes associated with these clusters were also suggested^88^.

Given this, we examined the composition of 45S rDNA repeats as well as their transcription. While the large and highly repetitive 45S rDNA clusters are not part of the genome assembly, it is possible to measure the copy number of *A. thaliana* and *A. arenosa* 45S rRNA genes using sequencing coverage (see Methods), and we find three accessions to have experienced massive loss of the *A. thaliana* rDNA loci (Fig. 2a), which we confirmed for one of the accessions (“AS90a”) by FISH analysis (Fig. 2b,c). However, there is massive copy number variation for 45S rRNA genes in *A. suecica* (Fig. 2a), and some accessions (e.g., the reference accession “ASS3”) have higher *A. thaliana* than *A. arenosa* 45S rRNA copy number (Fig. 2d,e).

**Figure 2.**
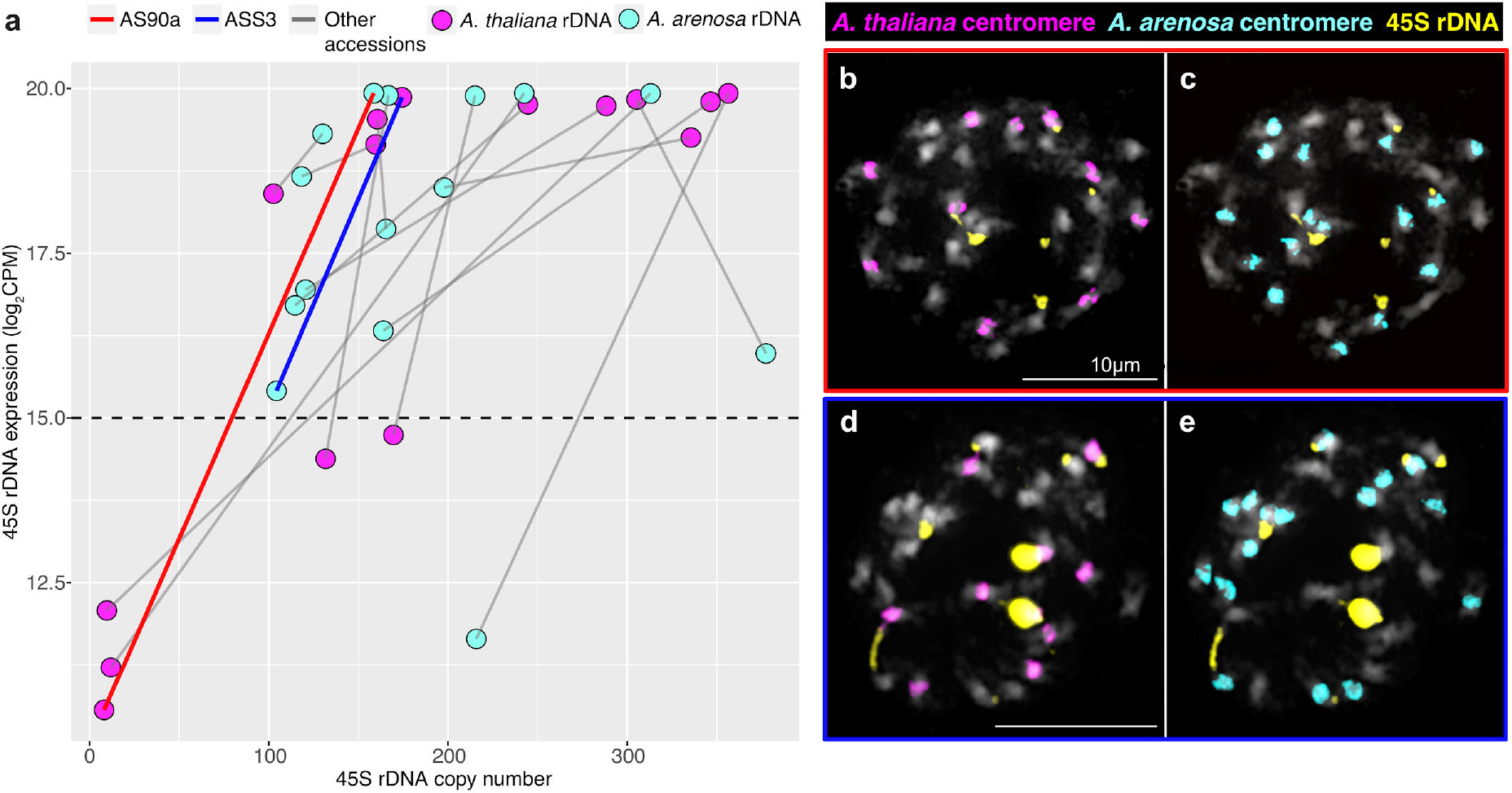
Expression and copy number variation of 45S rDNA in *A. suecica*. **a** The relationship between expression levels (log2 CPM) and copy number of 45S rDNA shows extensive variation of 45S rDNA copy number and varying direction of “nucleolar dominance”. Grey lines connect subgenomes of the same accession. Values above the dashed line are taken as evidence for the expression of a particular 45S rDNA allele, as this is above the maximum level of mis-mapping seen in the ancestral species here used as a control (see Supplementary Figure 6b). **b** and **c** FISH results of a natural *A. suecica* accession AS90a that has largely lost the rDNA cluster of the *A.thaliana* subgenome (8 copies calculated for the *A. thaliana* 45S rDNA and 159 copies of the *A. arenosa* 45S rDNA). **d** and **e** FISH result of a natural accession ASS3 that has maintained both ancestral rDNA loci (174 copies calculated for the *A. thaliana* 45S rDNA and 104 copies of the *A. arenosa* 45S rDNA).

Turning to expression, we also find nucleolar dominance to be variable in *A. suecica* (see Methods and Supplementary Fig. 6), with the majority of accessions expressing both 45S rRNA alleles, five exclusively expressing *A. arenosa* 45S rRNA, and one exclusively expressing *A. thaliana* 45S rRNA (Fig. 2a).

This extensive variation in 45S cluster size and expression is reminiscent of the genetically controlled intraspecific variation seen in *A. thaliana* (where different accessions express either the chromosome 2 or chromosome 4 rDNA cluster, or both^89, 90^), and is in agreement with a previous observation made in natural *A. suecica* that both rDNA clusters can be expressed^91^. This suggests that the phenomenon of nucleolar dominance can at least partly be explained by retained ancestral variation. However, the large-scale decrease in rDNA cluster size observed in some accessions may be a direct consequence of allopolyploidization itself, as synthetic *A. suecica* sometimes shows immediate loss of 45S rDNA (even as early as the F1 stage) and this too varies between siblings and generations (Supplementary Fig. 6a). Elimination of rDNA loci has also been previously observed in synthetic wheat^92^, and loss of rDNA sites has been reported at higher ploidy levels in strawberry^93^.

### 3. No evidence for abnormal transposon activity

The possibility that hybridization and polyploidization leads to a “genome shock” in the form of increased transposon activity has been much discussed^27, 28, 94, 95^. Evidence for TE proliferation following hybridization has been found for *Ty3/gypsy* retrotransposons in hybrid sunflower species^96^, though notably the hybrid sunflower species occupy habitats that are abiotically extreme^97^ which is also implicated in LTR proliferation^98^. On the other hand, analysis of TE expression in F1 hybrids between *A. thaliana* and *A. lyrata* found strong correlation, even under drought stress, to the parent species, as well as little alteration of the chromatin marks H3K9me2 and H3K27me3^99^ — although it remains unclear whether the F1 generation is not too early to study TE misregulation. Here we examine TE dynamics in natural *A. suecica*.

The two subgenomes of *A. suecica* differ massively in transposon content: there are almost twice as many annotated transposons in the *A. arenosa* as in the *A. thaliana* subgenome (66,722 vs 33,420; see Supplementary Figs. 5a and 7), and the true difference is almost certainly greater given that the *A. arenosa* subgenome assembly is less complete (and many of the missing regions are likely to be repeat-rich) and that the transposon annotation is biased towards *A. thaliana.* Has the combination of two such different genomes lead to increased transposon activity?

Our assembled *A. thaliana* subgenome does contain roughly 3,000 more annotated transposons than the TAIR10 *A. thaliana* reference genome, but this could reflect greater transposon number in the *A. thaliana* ancestors of this genome rather than increased transposon activity in *A. suecica.* In order to gain insight into transposon activity in *A. suecica*, we need to identify jumps that occurred after the species separated (and are thus only found in this species). We used the software PopoolationTE2^100^ to call presence-absence variation on a population-scale level using genome re-sequencing datasets for 15 natural *A. suecica* accessions, 18 *A. thaliana* accessions genetically close to *A. suecica*, and 9 *A. arenosa* lines. Of the 24,569 insertion polymorphisms called with respect to the *A. thaliana* subgenome, 8,767 were shared between *A. thaliana* and *A. suecica*, 7,196 were only found in *A. thaliana*, and 8,606 were only found in *A. suecica*. Of the 115,336 insertions on the *A. arenosa* subgenome of *A. suecica,* 13,177 were shared with *A. arenosa*, 83,964 were unique to *A. arenosa*, and 18,195 were unique to *A. suecica* (Supplementary Data 1a,b; Supplementary Figs. 8,9). Considering the number of transposons per individual genome (Fig. 3a), we see that most transposon insertions in a typical *A. thaliana* subgenome are also found in *A. thaliana*, and that the slightly higher transposon load in the *A. thaliana* subgenome is mainly due to these. The reason for this is likely a population bottleneck. In contrast, the number of recent insertions (that are unique to the species) is not higher in the *A. thaliana* subgenome, suggesting that transposon activity in this subgenome is not increased.

**Figure 3.**
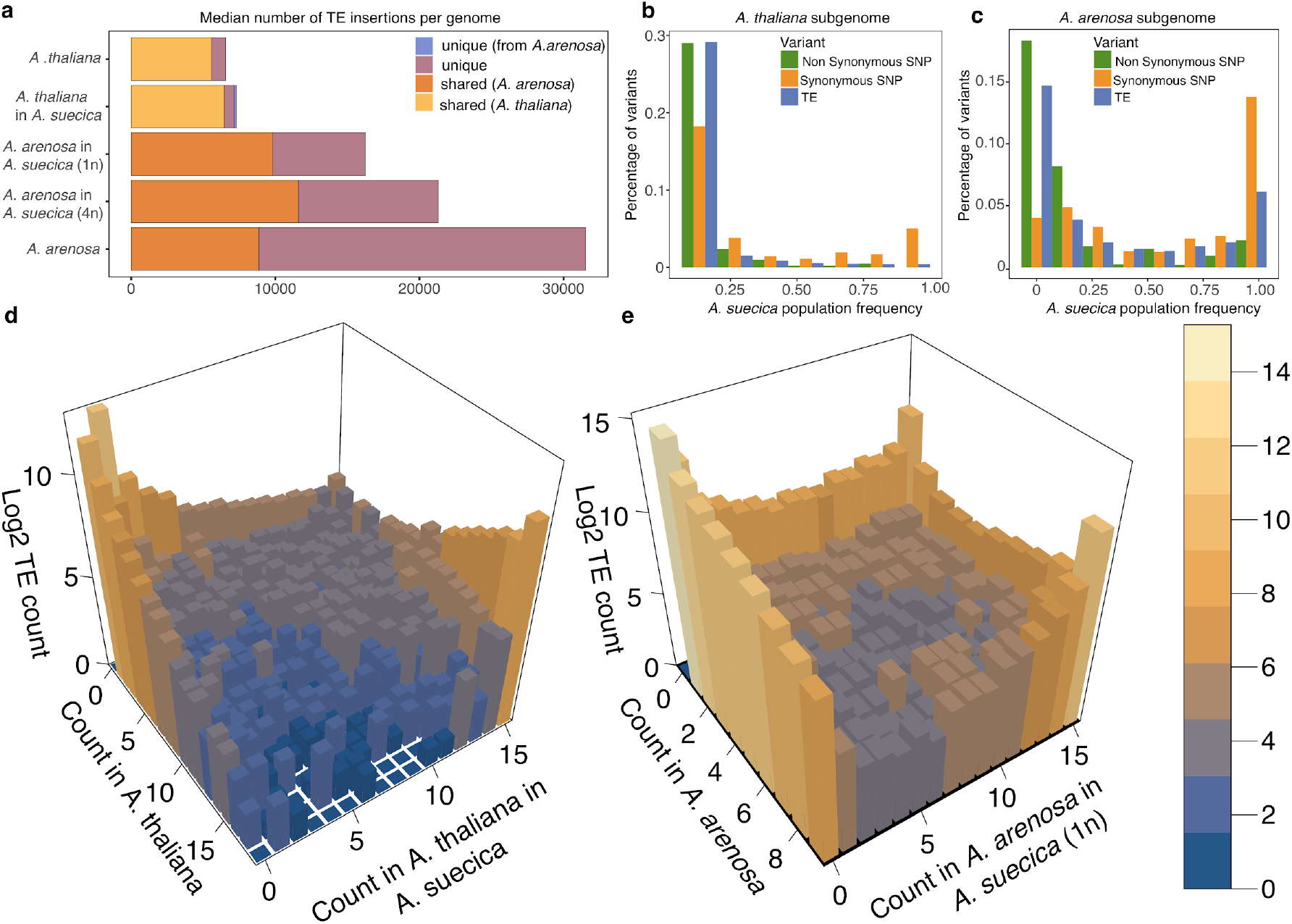
TE dynamics in *A. suecica* reveal no evidence for abnormal transposon activity. **a** Median TE insertions per genome. As the *A. arenosa* population is an autotetraploid outcrosser, 4 randomly chosen haploid *A. arenosa* subgenomes of *A. suecica* were combined to make a 4n *A. suecica*. *A. suecica* does not show an increase in private TE insertions compared with the ancestral species for both subgenomes, and shared TEs constitute a higher fraction of TEs in *A. suecica* reflecting the strong population bottleneck at its origin. Site-frequency spectra of non-synonymous SNPs, synonymous SNPs and TEs in the **b** *A. thaliana* and **c** *A. arenosa* subgenomes of *A. suecica* suggest that TEs are under purifying selection on both subgenomes. **d** 3D histogram of a joint TE frequency spectrum for *A. thaliana* on the x-axis and the *A. thaliana* subgenome of *A. suecica* on the y-axis **e** 3D histogram of a joint TE frequency spectrum for *A. arenosa* on the x-axis and the *A. arenosa* subgenome of *A. suecica* on the y-axis. **d** and **e** show stable dynamics of private TEs in *A. suecica* and a bottleneck effect on the ancestral TEs (shared) at the origin of the *A. suecica* species.

Turning to the *A. arenosa* subgenome, we see that a typical *A. suecica* contains only about half the number of transposons of a typical *A. arenosa* individual (Fig. 3a). However, the latter is an outcrossing tetraploid, and it is thus fairer to compare with the number of transposons in four randomly chosen *A. arenosa* subgenomes of *A. suecica* (shown as “*A. arenosa* in *A. suecica* (4n)” in Fig. 3a). This largely accounts for the observed difference, but there are still clearly fewer transposons in *A. suecica*. A population bottleneck likely explains much of this, but it is impossible to rule out a contribution of decreased transposon activity in *A. suecica* as well, which might be explained by its transition to self-fertilization, which is often associated with reduced TE activity^101^.

To sum up, we see no evidence for a burst of transposon activity accompanying polyploidization in *A. suecica*, a conclusion also supported by a lack of increase in transposon expression for both synthetic and natural *A. suecica* compared to the *A. thaliana* and *A. arenosa* on both subgenomes (Supplementary Fig. 9), in agreement with observations made in *A. thaliana* and *A. lyrata* F1 hybrids^99^. We do see clear traces of the population bottleneck accompanying the origin of *A. suecica*, however. The frequency distribution of polymorphic transposon insertions private in *A. suecica* is heavily skewed towards zero — almost certainly because of purifying selection because the distribution is more similar to that of non-synonymous SNPs than to that of synonymous SNPs (Fig. 3b,c). However, for both subgenomes, *A. suecica* also contains a large number of fixed or nearly-fixed insertions that are present in the ancestral species at lower frequency (Fig. 3d,e). These are likely to have reached high-frequency as a result of a bottleneck. Shared transposons are enriched in the pericentromeric regions of the genome depleted of protein-coding genes, while unique transposons insertions, which are generally at low frequency, show a more uniform distribution across the genome, consistent with evidence for stronger selection against transposon insertion in the relatively gene-dense chromosome arms^102, 103^ (Supplementary Fig. 10).

An interesting subset of recent transposon insertions unique to *A. suecica* are those that have jumped between the two subgenomes. We searched for full-length transposon copies that are present in both subgenomes of *A. suecica* and then assigned the resulting consensus sequences to either the *A. thaliana* or the *A. arenosa* ancestral genome using BLAST (see Methods). We were able to assign 15 and 56 consensus sequences as being specific to the *A. thaliana* and *A. arenosa* ancestral genome, respectively. Using these sequences, we searched our transposon polymorphism data for corresponding polymorphisms, and identified 1,515 *A. arenosa* transposon polymorphisms on the *A. thaliana* subgenome, and 496 *A. thaliana* transposon polymorphisms on the *A. arenosa* subgenome. Like other private polymorphisms, these are skewed towards rare frequencies, and are uniformly distributed across the (sub-)genome. Most of the transposons that have jumped into the *A. thaliana* subgenome are helitrons and LTR elements (Supplementary Fig. 12). LTR (copia) elements also make up most of the *A. thaliana* transposons segregating in the *A. arenosa* subgenome. The fact that roughly three times as many new insertions appear to have resulted from jumps from *A. arenosa* to *A. thaliana* than the other way around is notable. It is suggestive of higher transposon activity in the *A. arenosa* subgenome, but we have to consider differences in genome size and transposon number. If there were no differences in activity, we would expect the number of cross-subgenome jumps to be proportional to the number of potential source elements and the size of the target genome. As we have seen, the *A. arenosa* subgenome contains roughly twice as many transposons as the *A. thaliana* subgenome, but is about 20% larger. We would thus expect a 1.7-fold difference, not a three-fold one.

In conclusion, transposon activity in *A. suecica* appears to be governed largely by the same processes that governed it in the ancestral species.

### 4. No global dominance in expression between the subgenomes

Over time the traces of polyploidy are erased through an evolutionary process involving gene loss, often referred to as fractionation or re-diploidization^104–108^. Analyses of retained homeologs in ancient allopolyploids such as *A. thaliana*^109^, maize^55^, *B. rapa*^54^ and *Gossypium raimondii*^110^ have revealed that one “dominant” subgenome remains more intact, with more highly expressed homeologs compared to the “submissive” genome(s)^109^. This pattern of “biased fractionation” has not been observed in ancient autopolyploids^111^, such as pear^112^, and is believed to be allopolyploid-specific.

Studying genome expression dominance in contemporary allopolyploids is useful for understanding or predicting which of the subgenomes will likely be refractory to, and which will likely experience this fractionation process more, over time^55^. Subgenome dominance in expression has been reported for a number of more recent allopolyploids such as strawberry^6^, peanut^8^, *Spartina*^68^*, T. miscellus*^113^, monkeyflower^17^ and synthetic *B. napus*^114^. However, some allopolyploids display even subgenome expression, among them *C. bursa-pastoris*^10, 12^, white clover^13^, *A. kamachatica*^70^ and *B. hybridum*^14^.

Subgenome dominance is often linked to differences in transposon content^6^ and/or large genetic differences between subgenomes^115^. This makes *A. suecica*, with 6 Mya divergence between the gene-dense *A. thaliana* and the transposon-rich *A. arenosa*, a promising candidate to study this phenomenon at unprecedented resolution. Previous reports on subgenome dominance in *A. suecica* are conflicting, suggesting a bias to either the *A. thaliana*^116^ or the *A. arenosa*^117^ subgenome.

To investigate the evolution of gene expression in *A. suecica*, we generated RNA-seq data for 15 natural *A. suecica* accessions, 15 closely related *A. thaliana* accessions, 4 *A. arenosa* individuals, a synthetically generated *A. suecica* from a lab cross (the 2nd and 3rd hybrid generations) and the parental lines of this cross. Each sample had 2-3 biological replicates (Supplementary Data 2). On average, we obtained 10.6 million raw reads per replicate, of which 7.6 million reads were uniquely mapped to the *A. suecica* reference genome and 14,041 homeologous gene pairs (see Methods, Supplementary Fig. 13).

Considering the difference in expression between homeologous genes, we found no general bias towards one or the other subgenome of *A. suecica*, for any sample or tissue, including synthetic *A. suecica* (Fig. 4a and Supplementary Fig. 14a). This strongly suggests that the expression differences between the subgenomes have not changed systematically through polyploidization, and is in contrast to previous studies, which reported a bias towards the *A. thaliana*^116^ or the *A. arenosa*^117^ subgenome, likely because RNA-seq reads were not mapped to an appropriate reference genome.

**Figure 4.**
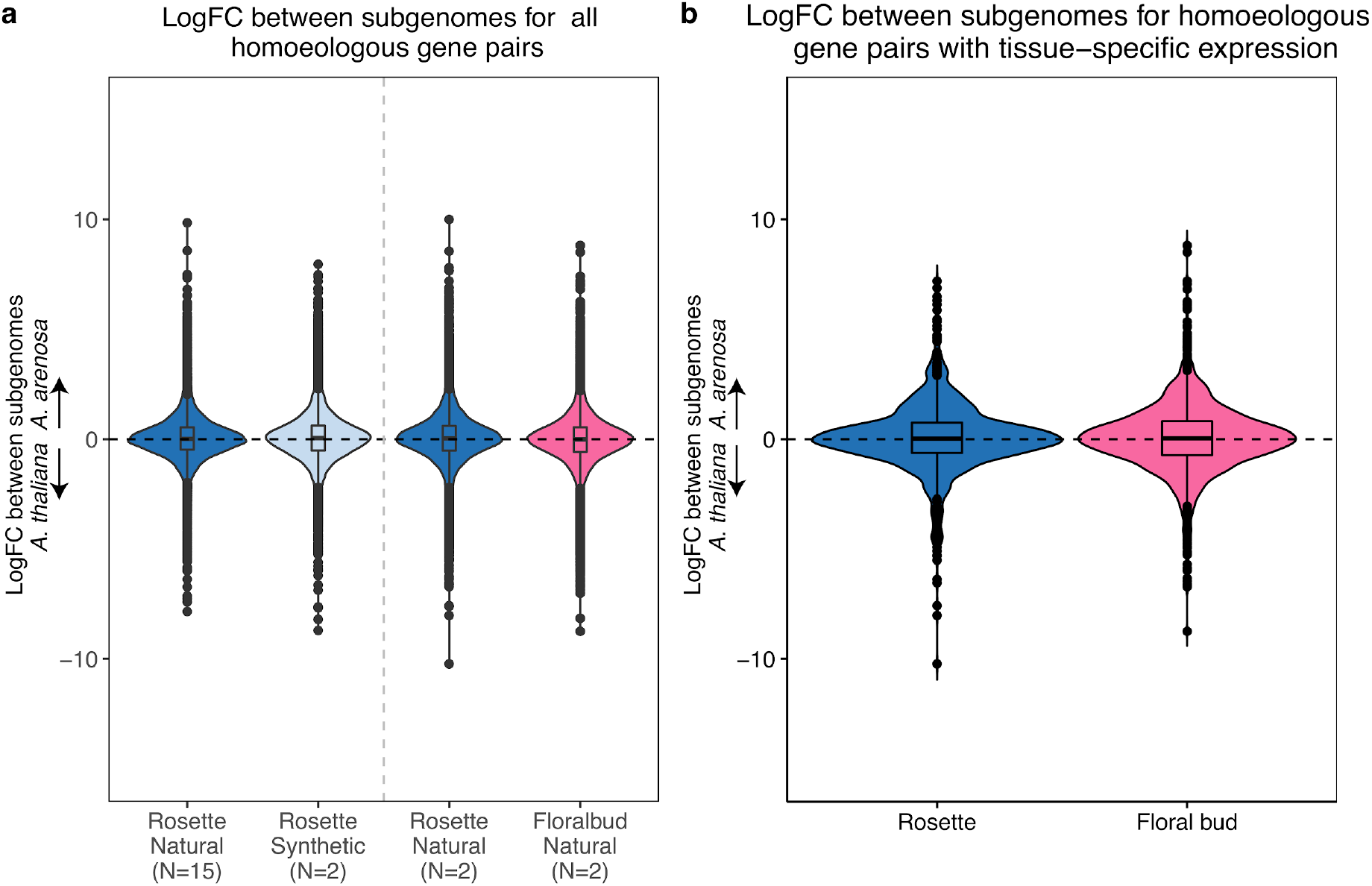
Patterns of gene expression between the subgenomes of *A. suecica* in rosettes and floral buds. **a** Violin plots of the mean log fold-change between the subgenomes for the 15 natural *A. suecica* accessions and two synthetic lines for whole rosettes. Mean log fold-change for the two accessions (“ASS3” and “AS530”) where transcriptome data for both whole rosettes and flower buds were available. All the distributions are centered around zero suggesting even subgenome expression. **b** Violin plots for the mean log fold-change between the subgenomes for genes with tissue-specific expression. At least one gene in a homeologous gene pair was required to show tissue-specific expression.

The set of genes that show large expression differences between the subgenomes appears not to be biased towards any particular gene ontology (GO) category, and is furthermore not consistent between accessions and individuals (Fig. 4b, Supplementary Fig. 14b,c). This suggests that many large subgenome expression differences are due to genetic polymorphisms within *A. suecica* rather than fixed differences relative to the ancestral species. Levels of expression dominance were reported to vary across tissues in natural *C. bursa-pastoris*^11^ and also resynthesized cotton^118^. To test whether expression dominance can vary for tissue-specific genes, we examined homeologous gene-pairs where at least one gene in the gene pair showed tissue specific expression, in whole-rosettes and floral buds. We do not find evidence for dominance between subgenomes in tissue specific expression either (Fig. 4b). Interestingly, the 897 genes with significant expression in whole rosettes for both homeologs showed GO overrepresentation that included both photosynthesis and chloroplast related functions (Supplementary Table 1). This result suggests that the *A. arenosa* subgenome has established important cyto-nuclear communication with the chloroplast inherited from *A. thaliana*, rather than being silenced. 2,176 gene pairs with floral bud specific expression for both homeologs were overrepresented for GO terms related to responses to chemical stimuli, such as auxin and jasmonic acid, which may reflect early developmental changes in this young tissue (Supplementary Table 1). Although flowers of selfing *A. thaliana* and *A. suecica* are scentless and are much smaller than those of the outcrosser *A. arenosa*^72^, this result suggests the “selfing syndrome”^119^ has not hugely impacted the transcriptome of floral buds in *A. suecica*, at least at this stage of development.

In summary, we find no evidence that one subgenome is dominant and contributes more to the functioning of *A. suecica*. On the contrary, homeologous gene pairs are strongly correlated in expression across tissues.

### 5. Evolving gene expression in *A. suecica*

The previous section focused on differences in expression between the subgenomes, between homeologous copies of the same gene within the same individual. This section will focus on differences between individuals, between homologous copies of genes that are part of the same (sub-)genome. To provide an overview of expression differences between individuals we performed a principal component analysis (PCA) on gene expression separately for each (sub-)genome. For both subgenomes, the first principal component separates *A. suecica* from the ancestral species and the synthetic hybrid (Fig. 5a,b, Supplementary Fig. 15), suggesting that hybridization does not automatically result in large-scale transcriptional changes, and that altered gene expression changes in natural *A. suecica* have evolved over time. Given the limited time involved, and the fact the genes that have changed expression are far from random with respect to function (Fig. 5c), we suggest that the first principal component primarily captures trans-regulated expression changes in *A. suecica* that are likely adaptive.

**Figure 5.**
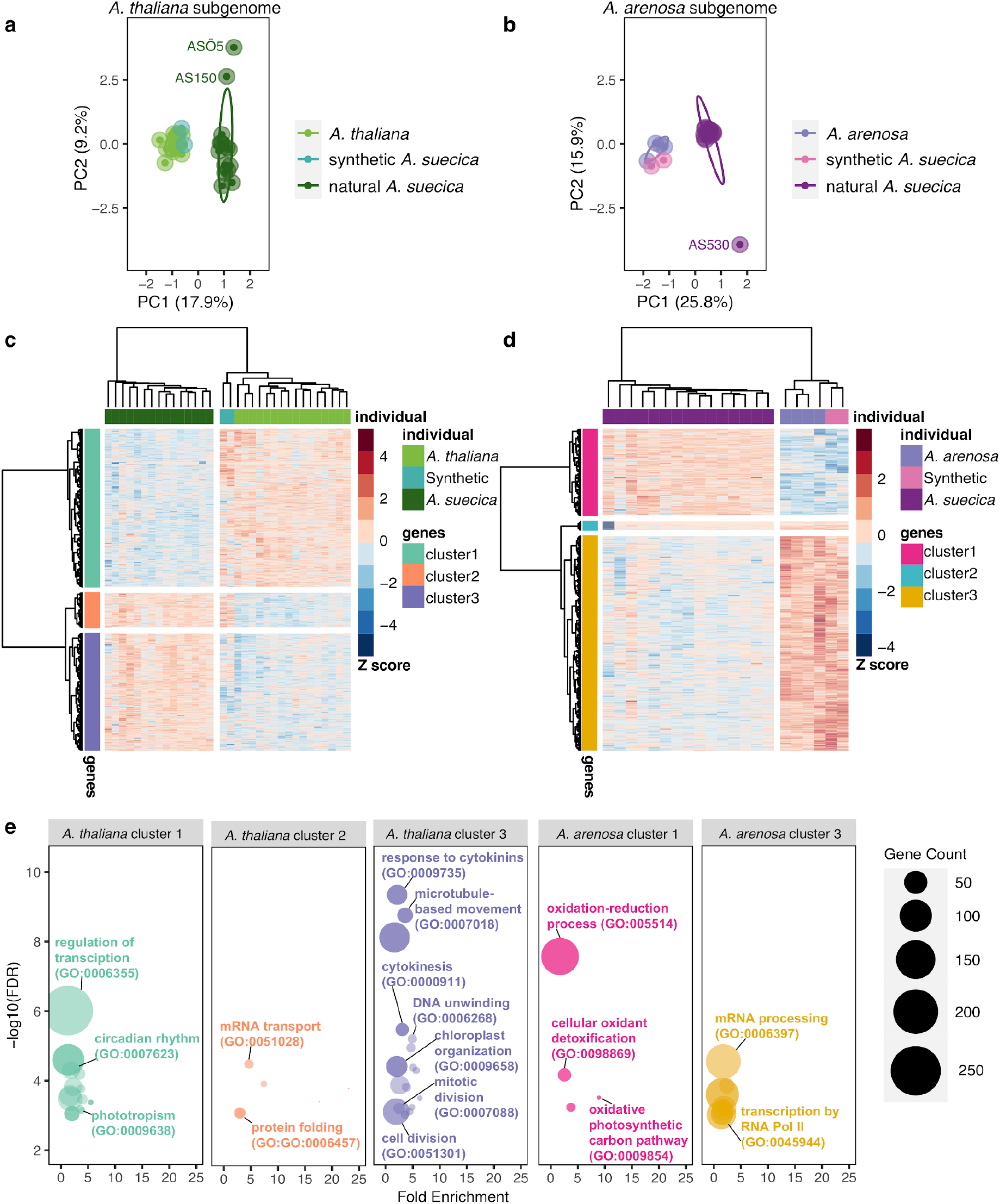
Differential gene expression analysis in *A. suecica*. Patterns of differential gene expression in *A. suecica* support adaptation to the whole-genome duplication for the *A. thaliana* subgenome and adaptation to the new plastid environment for the *A. arenosa* subgenome. **a** PCA for *A. thaliana*, *A. thaliana* subgenome of natural and synthetic *A. suecica* lines. PC1 separates natural *A. suecica* from the ancestral species and the synthetic lines. **b** PCA for *A. arenosa*, *A. arenosa* subgenome of natural and synthetic *A. suecica* lines. PC1 separates natural *A. suecica* from the ancestral species and the synthetic lines, whereas PC2 identifies outlier accessions discussed further below (see Fig. 6). **c, d** Heatmap of differentially expressed genes (DEGs) for the two subgenomes of *A. suecica*. Positive numbers (red color) indicate higher expression. Genes and individuals have been clustered based on similarity in expression, resulting in clusters discussed in the text. **e** Gene ontology enrichment for each cluster in **c** and **d**. Categories discussed in the text are highlighted.

To further characterize expression changes in natural *A. suecica* we analyzed differentially expressed genes (DEGs) on both subgenomes compared to the corresponding ancestral species. The total number of DEGs was 4,186 and 4,571 genes for the *A. thaliana* and *A. arenosa* subgenomes, respectively (see Methods, Supplementary Data 2). These genes were clustered based on the pattern of change across individuals (Fig. 5c,d) and GO enrichment analysis was carried out for each cluster (Fig. 5e, Supplementary Table 2).

For the *A. thaliana* subgenome, we identified three clusters. Cluster 1 comprised 2,135 genes that showed decreased expression in *A. suecica* compared to *A. thaliana*. These genes are strongly enriched for transcriptional regulation, which may be expected as we are examining DEGs between the species. Also notable are enrichments for circadian rhythm function and phototropism, which may be related to the ecology of *A. suecica* and its post-glacial migration to the Fennoscandinavia region (Fig. 1a).

Cluster 2 consisted of 468 genes that are over-expressed in both natural and synthetic *A. suecica* relative to *A. thaliana*. These expression changes are thus most likely an immediate consequence of hybridization presumably reflecting trans-regulation. Genes in this cluster are enriched for “mRNA transport” and “protein folding”. The importance of the adjustment of protein homeostasis has been reported previously in experimentally evolved stable polyploid yeast^120^. Notably, the synthetic lines used in the expression analysis were selected to be healthy-looking, and did not show signs of aneuploidy (Supplementary Fig. 17).

Cluster 3 consisted of 1,583 genes that show increased expression in *A. suecica* compared to *A. thaliana*, and several of the enriched GO categories, such as microtubule-based movement, cytokinesis, meiosis and cell division, suggest that the *A. thaliana* subgenome of *A. suecica* is adapting to polyploidy at the level of basic cell biology. That there has been strong selection for this seems likely given that aneuploidy is frequent in synthetic *A. suecica* (Supplementary Fig. 16), while natural *A. suecica* has a stable and conserved karyotype. Importantly, there is independent evidence for adaptation to polyploidy via modifications of the meiotic machinery in the other ancestor of *A. suecica*, *A. arenosa*, as well^23, 121, 122^, although we see very little overlap in the genes involved (Supplementary Fig. 16). The nature of these changes in the *A. thaliana* subgenome of *A. suecica* will require further investigation, but we note that there is enrichment (see Methods, Supplementary Data 2) for Myb family transcription factor binding sites^123^ among upregulated genes in cluster 3.

For the *A. arenosa* subgenome, we also found three clusters of DEGs (Fig. 5d) with GO enrichment for two of them (Fig. 5e, Supplementary Table 2). Cluster 1 consisted of 1,278 genes that show increased expression in natural *A. suecica* compared to *A. arenosa* and synthetic *A. suecica,* and are enriched for plastid-related functions, including oxidation-reduction and the oxidative photosynthetic carbon pathway. We hypothesize that this may be due to selection on the *A. arenosa* subgenome to restore communication with the new plastid environment as plastid genomes were maternally inherited from *A. thaliana*. We also examined genes that show structural evidence for direct plastid-nuclear interactions in *A. thaliana* using CyMIRA^124^. Out of a total of 69 genes, 12 overlap genes identified in Cluster1, more than expected by chance (p-value 0.0072; one-sided Fisher Exact Test, one sided; Supplementary Data 2). Cluster 3 consists of 3,166 genes that show decreased gene expression in *A. suecica* compared to *A. arenosa* and synthetic *A. suecica*. These genes were primarily enriched for mRNA processing and epigenetic regulation of gene expression (Supplementary Table 2) and positive regulation of transcription by RNA polymerase II, which might suggests differences in the epigenetic regulation of expression between *A. arenosa* and *A. suecica*. Cluster 2 (127 genes), finally, did not have a GO overrepresentation and showed an intriguing pattern discussed in the next section.

### 6. Homeologous exchange contributes to variation in gene expression

The second principal component for gene expression identified three outlier-accessions of *A. suecica*, two for the *A. thaliana* subgenome (Fig. 5a) and one for the *A. arenosa* subgenome (Fig. 5b). While closely examining the latter accession, “AS530”, we realized that it is responsible for the cluster of genes with distinct expression patterns but no GO enrichment just mentioned (Fig. 5d, Cluster 2). Genes from this cluster were significantly downregulated on the *A. arenosa* subgenome (Fig. 6a) and upregulated on the *A. thaliana* subgenome (Fig. 6b) — for AS530 only. The further observation that 104 of the 127 genes (Supplementary Fig. 20a) in the cluster are located in close proximity in the genome, pointed to a structural rearrangement. The lack of DNA sequencing coverage on the *A. arenosa* subgenome around these 104 genes and the doubled coverage for their homeologs on the *A. thaliana* subgenome, strongly suggested a homeologous exchange (HE) event resulting in AS530 carrying four copies of the *A. thaliana* subgenome and zero copies of the *A. arenosa* genome with respect to this this, roughly 2.5 Mb region of the genome (Fig. 6c). This explanation was further supported by HiC data, which showed clear evidence for interchromosomal contacts between *A. thaliana* subgenome chromosome 1 and *A. arenosa* subgenome chromosome 6 around the breakpoints of the putative HE in AS530 (Fig. 6 d,e), and by multiple discordant Illumina paired-end reads at the breakpoints between the homeologous chromosomes, which independently support the HE event (Supplementary Fig. 19a-d).

**Figure 6.**
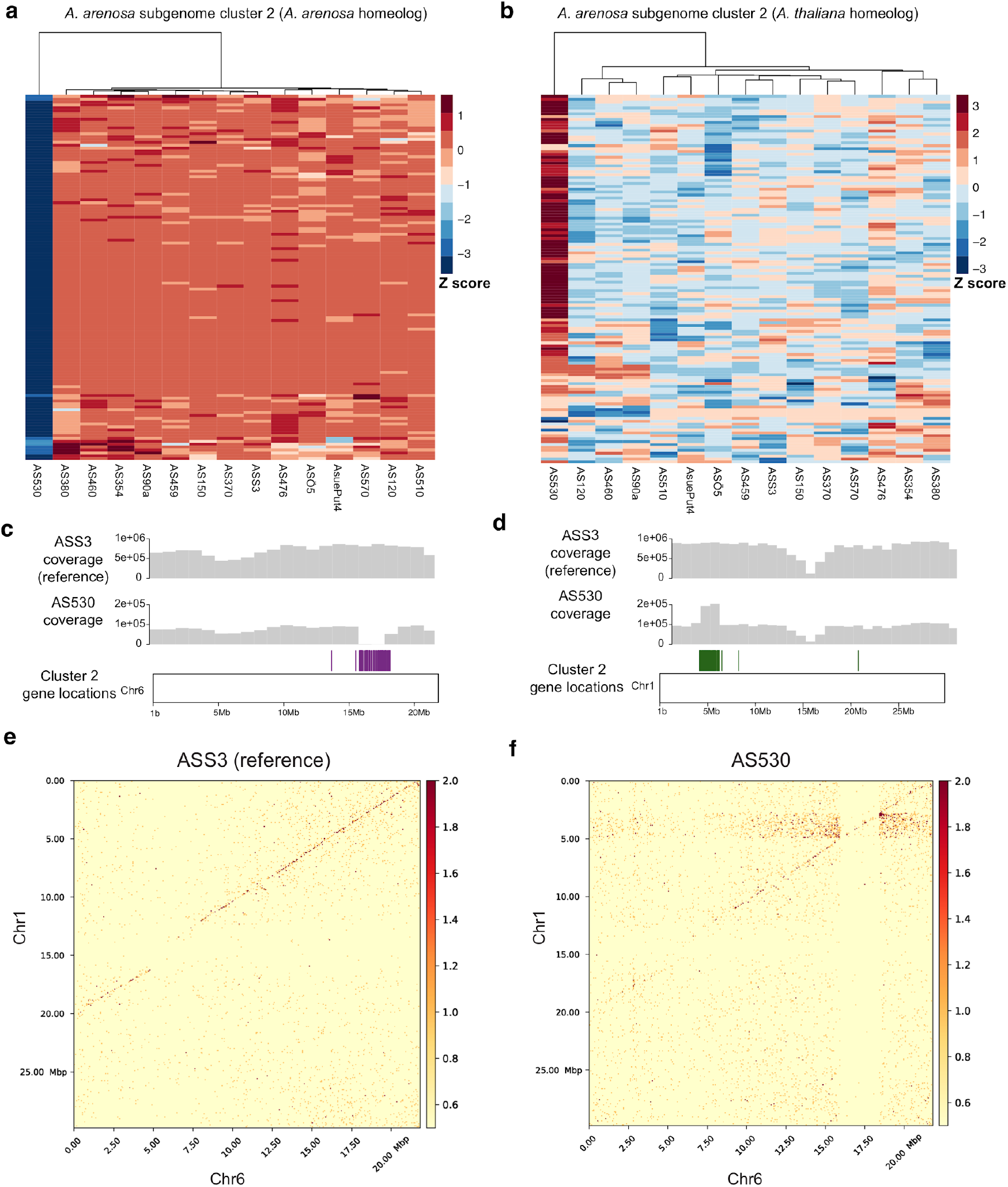
Homeologous exchange contributes to expression variance within *A. suecica*. **a** Cluster 2 of Fig. 5d explains the outlier accession AS530 which is not expressing a cluster of genes on the *A. arenosa* subgenome. **b** Homeologous genes of this cluster on the *A. thaliana* subgenome of *A. suecica* show the opposite pattern and are more highly expressed in AS530 compared to the rest of the population. **c** 97 of the 122 genes from cluster 3 are located in close proximity to each other on the reference genome but appear to be deleted in AS530 based on sequencing coverage. **d** The *A. thaliana* subgenome homeologs have twice the DNA coverage, suggesting they are duplicated. **e** HiC data show (spurious) interchromosomal contacts at 25 Kb resolution between chromosome 1 and chromosome 6 around the breakpoint of the cluster of 97 genes in AS530 but not in reference accession ASS3.

Based on this we examined the two outlier *A. suecica* accessions for the *A. thaliana* subgenome (Fig. 5a; “AS150” and “ASÖ5”), and found that they likely share a single HE event in the opposite direction (four copies of the *A. arenosa* subgenome and no copies of the *A. thaliana* subgenome for a region of roughly 1.2Mb in size, see Supplementary Figure 18). This demonstrates that HE occurs in *A. suecica* and contributes to the intraspecific variation we observed in gene expression (Fig 5a, b). HE in allopolyploids is a main source of diversity, causing phenotypic changes in flower color in synthetic polyploid peanut^9^ and extensive phenotypic change in synthetic polyploid rice at a population level^125^. However, the majority of HEs are probably deleterious as they will lead to gene loss: although the *A. thaliana* and *A. arenosa* genomes are largely syntenic, AS530 is missing 108 genes (Supplementary Figure 19) that are only present on the *A. arenosa* subgenome segment that has been replaced by the homeologous segment from the *A. thaliana* subgenome, and AS150/ASÖ5 are missing 53 genes that were only present on the *A. thaliana* subgenome.

## Conclusion

This study has focused on the process of polyploidization in a natural allotetraploid species, *A. suecica*, generated roughly 16 kya through the hybridization of two species, *A. thaliana* and *A. arenosa*, which differ substantially in everything from genome size and chromosome number to mating system and ecology. Our study is one of a growing number of studies focusing on natural rather than domesticated polyploid, but is unparalleled in its resolution thanks to one of the parents being a major model species.

Our main conclusion from this study is that polyploid speciation, at least in this case, appears to have been a gradual process rather than some kind of “event”. We confirmed previous results that genetic polymorphism is largely shared with the ancestral species, demonstrating that *A. suecica* did not originate through a single unique hybridization event, but rather through multiple crosses^20^. We also find no evidence for “genome shock” (i.e. major genomic changes linked to structural and functional changes) that has often been suggested to accompany polyploidization and hybridization. The genome has not been massively rearranged, transposable elements are not out of control, and there is no subgenome dominance in expression. On the contrary, we find evidence of genetic adaptation to “stable” life as a polyploid, in particular changes to the meiotic machinery and in interactions with the plastids. These findings made in natural *A. suecica*, together with the observation that experimentally generated *A. suecica* are often unviable and do exhibit evidence of genome rearrangements, similar to the young allopolyploid species in *Tragapogon* and monkeyflower, suggest that the most important bottleneck in polyploid speciation may be selective. If this is true, domesticated polyploids may not always be representative of natural polyploidization, because of human intervention. Darwin famously argued that “Natura non facit saltum”^126^ — we suggest that natural polyploids are no exception from this, but note that many more species will have to be studied before it is possible to draw general conclusions.

## Supplemental figures

**Supplementary Figure 1.**
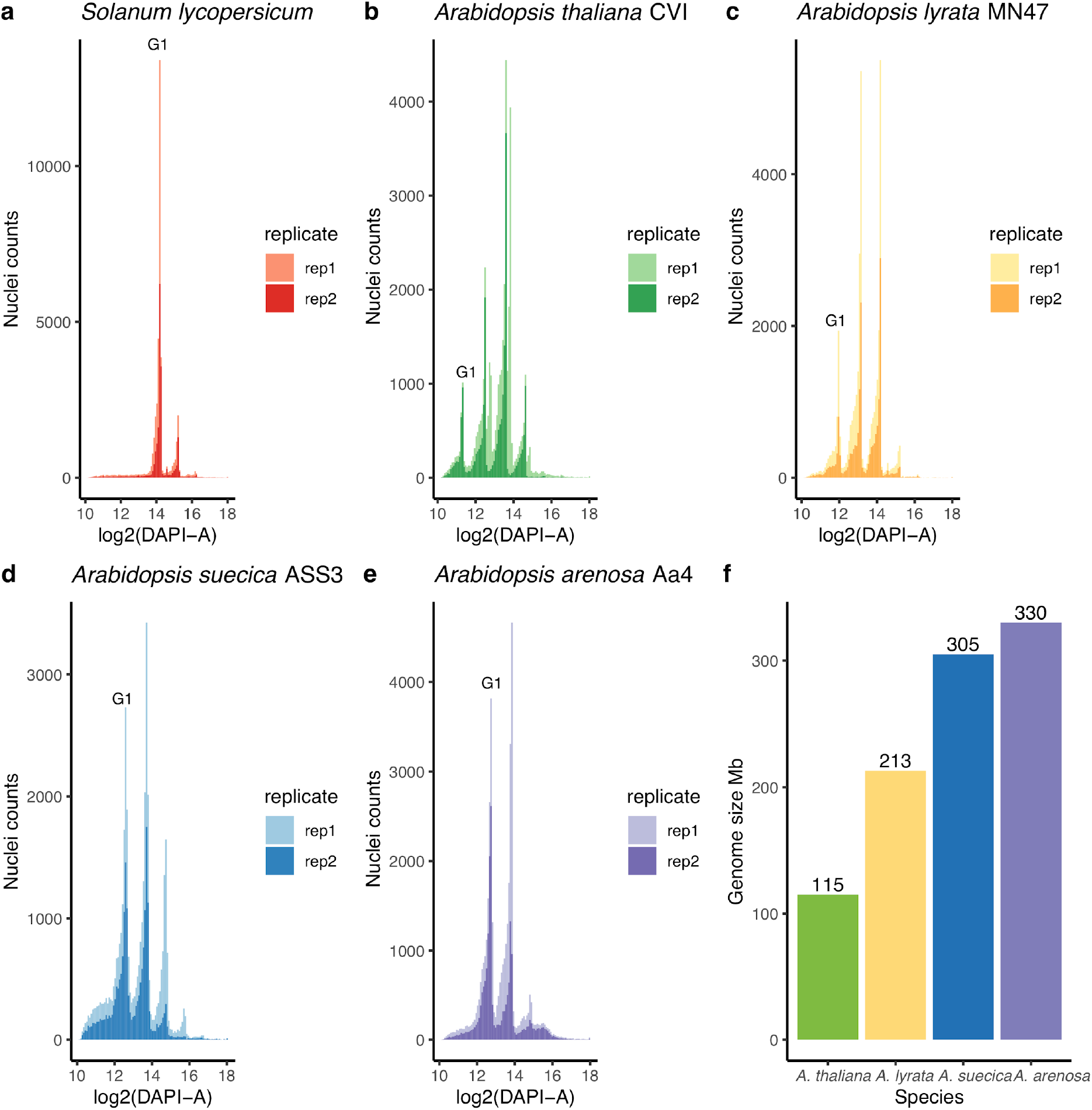
Measuring genome sizes of *Arabidopsis* species using flow cytometry. **a** FACs sorting of *Solanum lycopersicum* cells from 3 week old leaf tissue for two replicates. G1 represents the peak denoting the G1 phase of the cell cycle. Cells in the G1 phase have 2C DNA content (i.e. a 2N genome). **b** *A. thaliana* “CVI” accession **c** *A. lyrata* “MN47” (the reference accession) **d** *A. suecica* “ASS3” (the reference accession) **e** autopolyploid *A. arenosa* accession “Aa4” **f** Bar chart shows calculated genome sizes (rounded to the nearest whole number) for each species using *Solanum lycopersicum* as the standard.

**Supplementary Figure 2.**
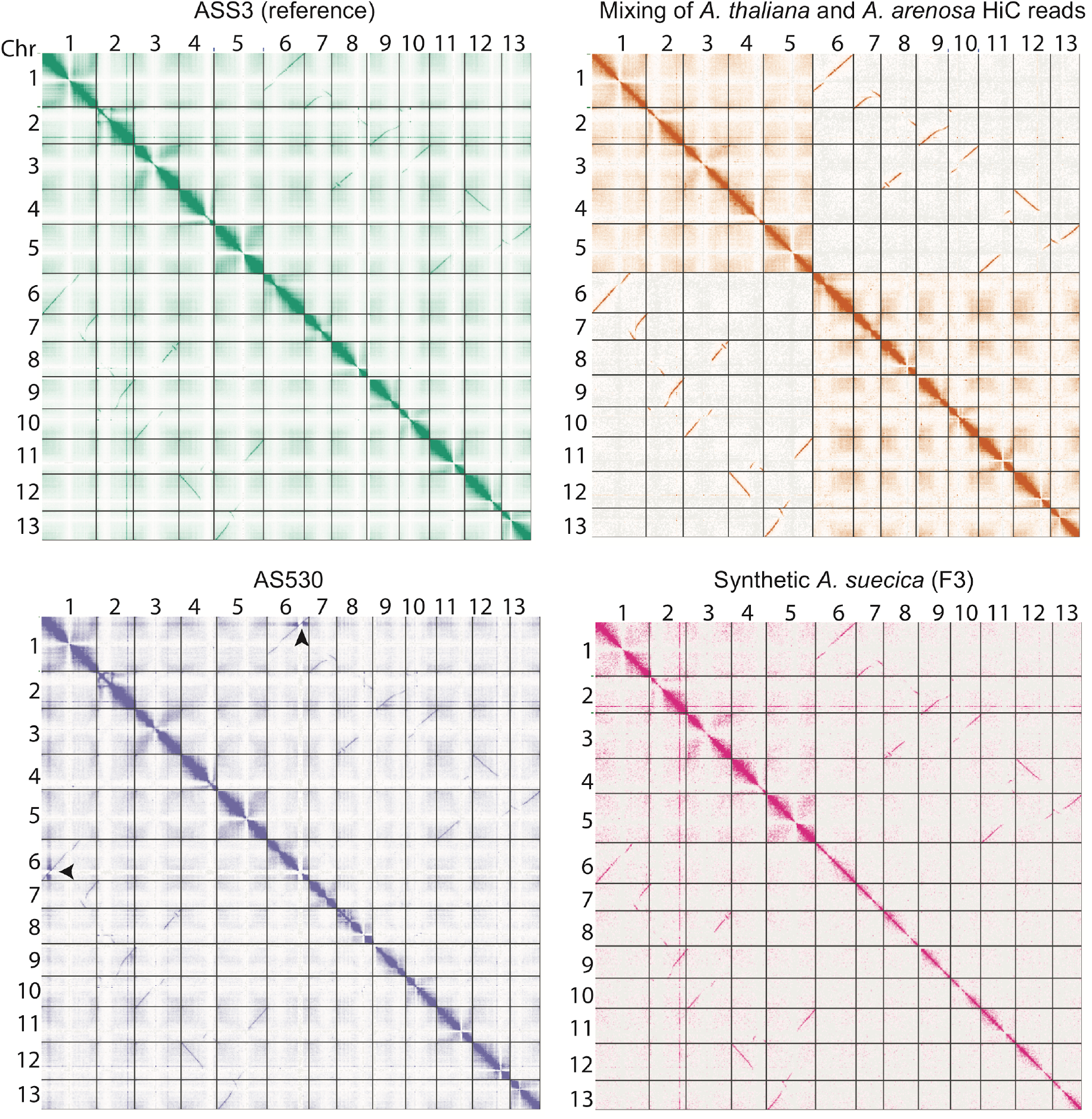
HiC as a tool to investigate structural rearrangements. **a** HiC contact map for the full chromosome-level genome assembly of *A. suecica*. **b** Mixing of *A. thaliana* and *A. arenosa* HiC reads suggest interchromosomal contacts between homeologous chromosomes is a result of mis-mapping for HiC reads. Such mis-mapping is typically filtered out in short read DNA and RNA datasets using insert size and proper pairs mapping filters, however in HiC long range chromosomal contacts are not filtered out. **c** Accession “AS530” with the region of homeologous exchange highlighted with an arrow (Figure 6), no other rearrangements were observed. **d** HiC of synthetic *A. suecica* (F3).

**Supplementary Figure 3.**
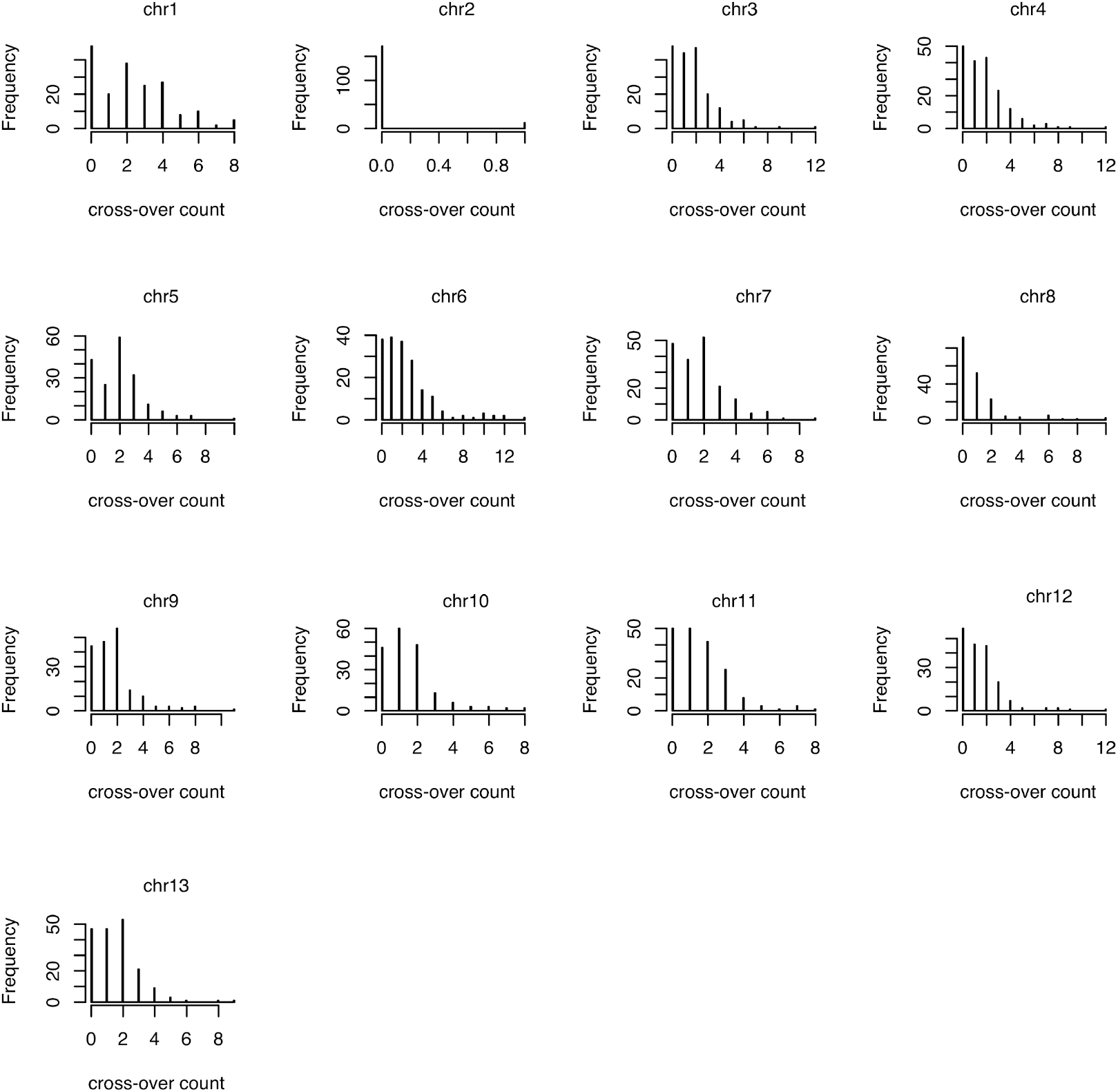
Crossover counts in an *A. suecica* F2 population. Per chromosome crossover counts in our F2 population (N=185). Chromosome 2 had too few SNPs to be analysed in our cross due to the recent bottleneck in *A. suecica*^20^.

**Supplementary Figure 4.**
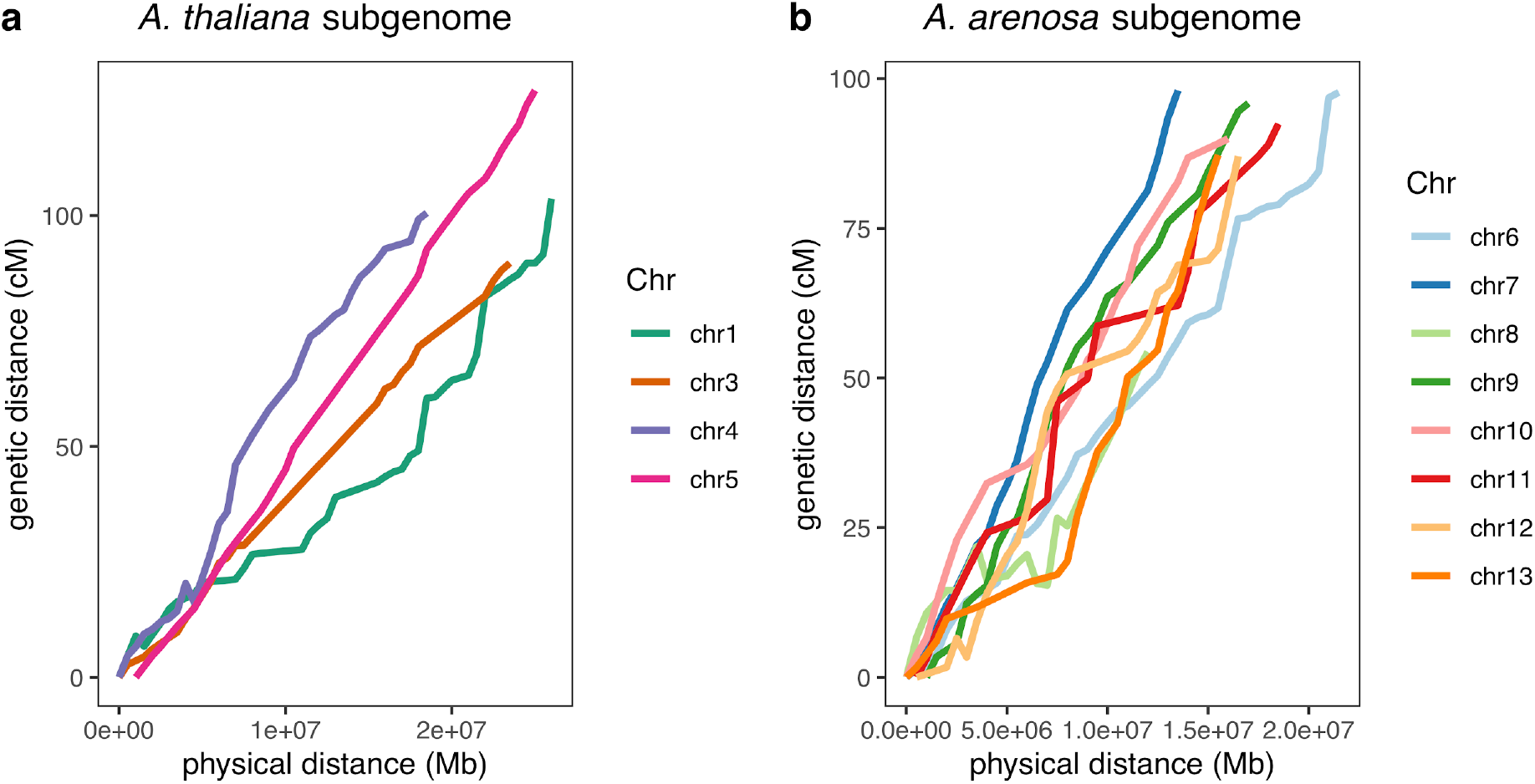
A genetic map for *A. suecica*. Physical distance (Mb) vs genetic distance (cM) is plotted for each: **a** *A. thaliana* subgenome and; **b** *A. arenosa* subgenome chromosome. Chromosome 2 is not plotted as there are too few SNPs on this chromosome in our cross, due to the recent bottleneck in *A. suecica*^20^

**Supplementary Figure 5.**
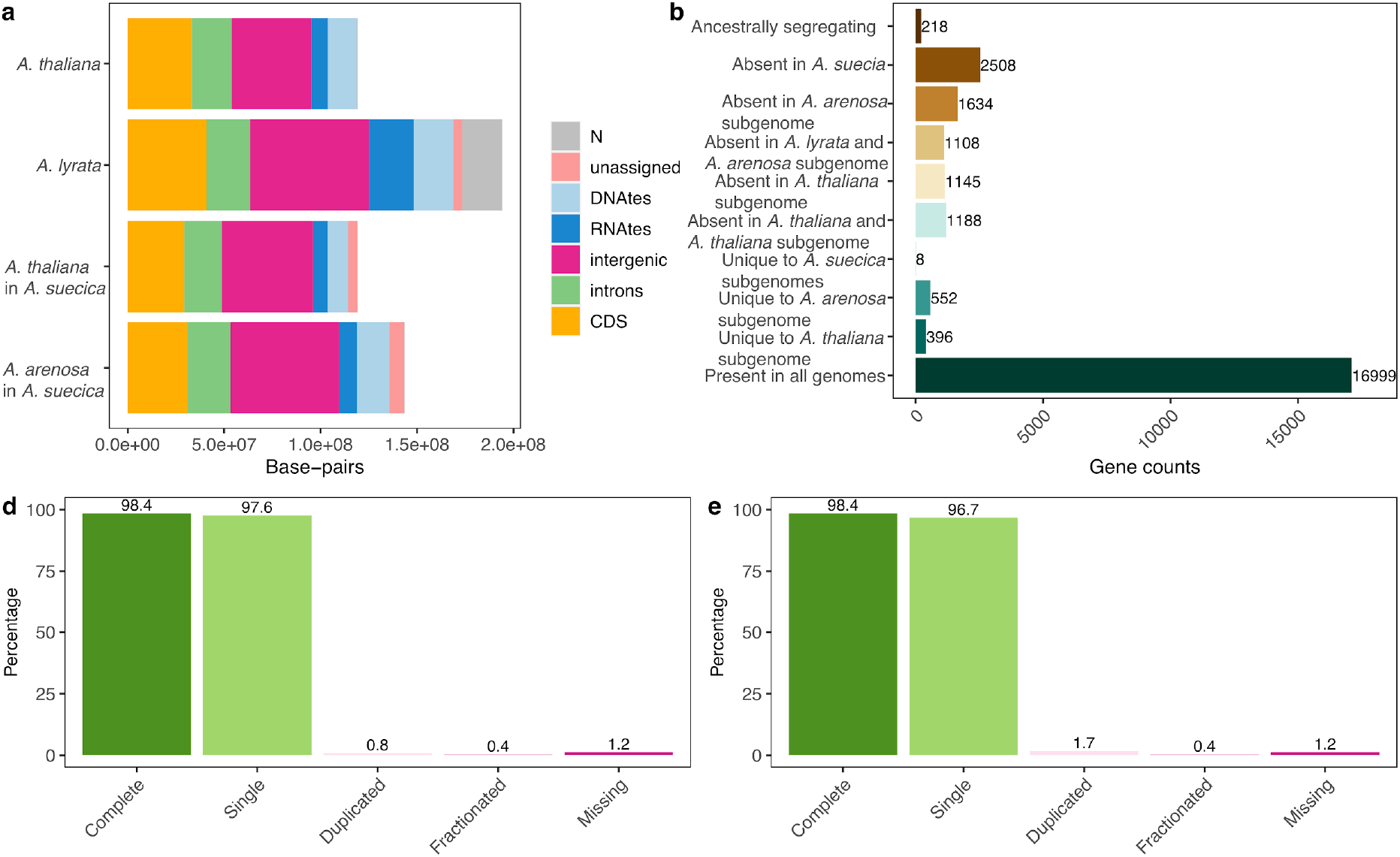
Genome composition and orthologous gene relationships in *A. suecica*. **a** Genome composition of the *A. suecica* subgenomes and the ancestral genomes of *A. thaliana* and *A.lyrata* (here a substitute reference for *A. arenosa* because it is annotated). **b** Counts of orthologous relationships between the subgenomes of the reference *A. suecica* genome and the reference *A. thaliana* and *A. lyrata* genome. Ancestrally segregating genes are genes that are shared between the *A. thaliana* reference and the *A. arenosa* subgenome or shared between the *A. lyrata* reference and the *A. thaliana* subgenome. Therefore they most likely represent genes ancestrally segregating in the ancestor of *A. thaliana* and *A. lyrata*. BUSCO analysis of *A. suecica* using the BUSCO set for eudicots for the **d** *A. thaliana* and **e** *A. arenosa* subgenome.

**Supplementary figure 6.**
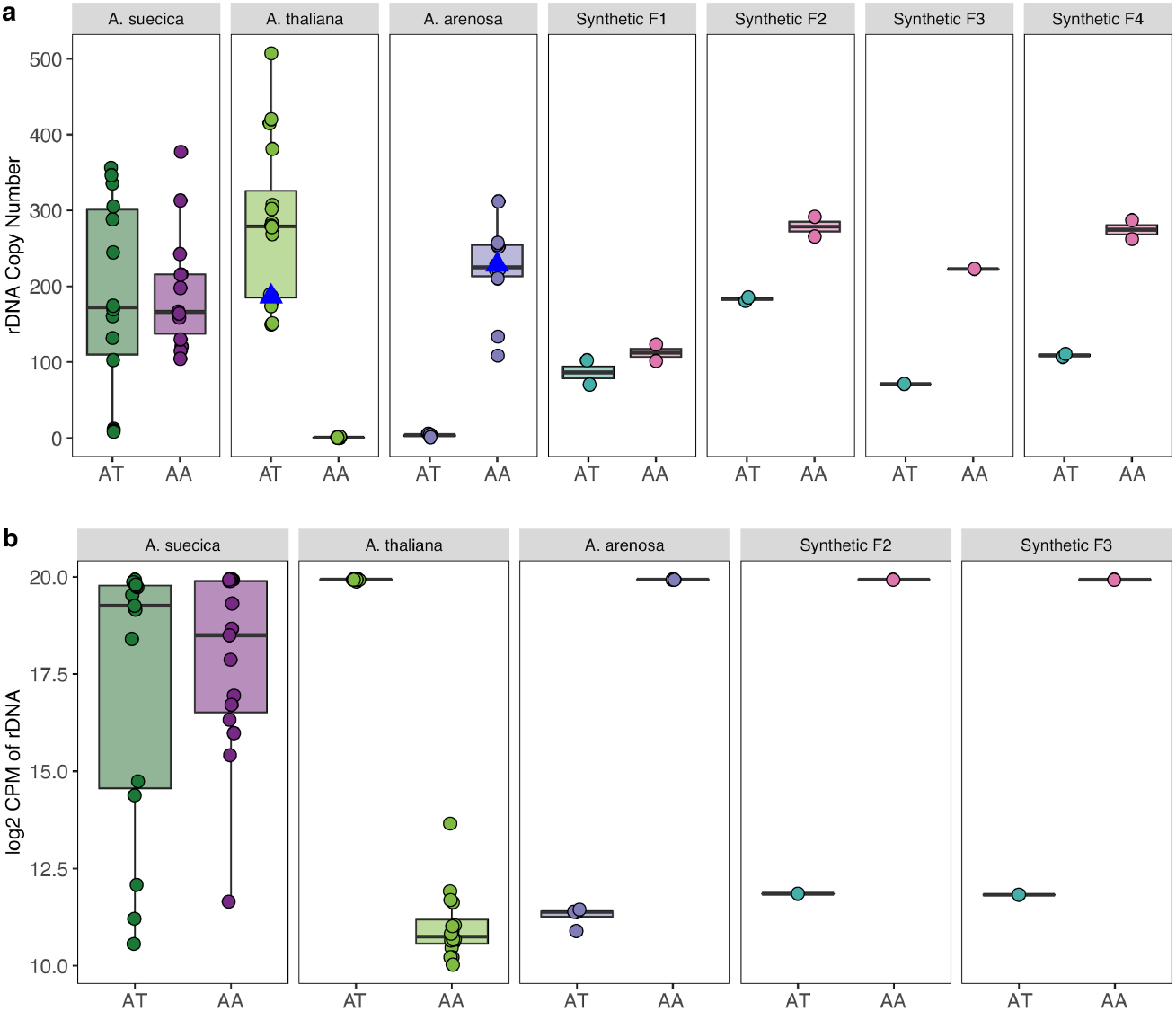
rDNA copy number variation and expression. **a** Copy number of *A. thaliana* and *A. arenosa* rDNA in natural *A. suecica*, ancestral species and synthetic lines. Blue triangles represent the *A. thaliana* and *A.arenosa* parent lines of the synthetic *A. suecica* cross. AT represents results when mapping to the *A. thaliana* consensus sequence and AA to the *A. arenosa* consensus sequences for the 45S rRNA **b** Expression (log2 CPM) of *A. thaliana* and *A. arenosa* rDNA in natural *A. suecica*, ancestral species and synthetic lines. Accessions with log2 CPM of >=15 was taken as evidence for expression for the *A. thaliana* and *A. arenosa* 45S rRNA in *A. suecica*, as this CPM value was above the maximum level of mis-mapping observed in the ancestral species (*A. thaliana* mapping to the *A. arenosa* 45S rRNA).

**Supplementary Figure 7.**
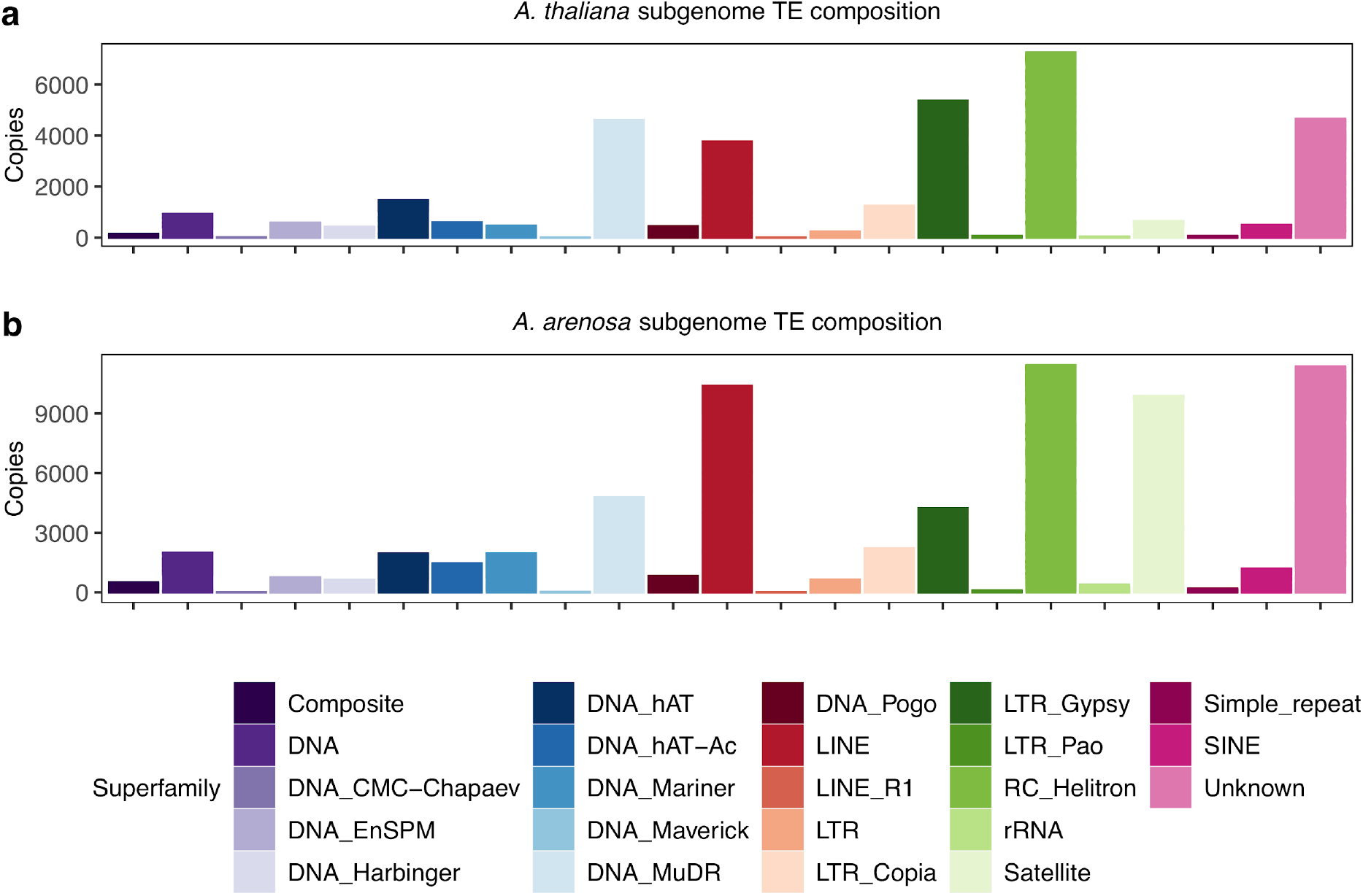
TE-composition of the *A. suecica* reference genome. TE composition of the **a** *A. thaliana* and **b** *A. arenosa* subgenome of *A. suecica*.

**Supplementary Figure 8.**
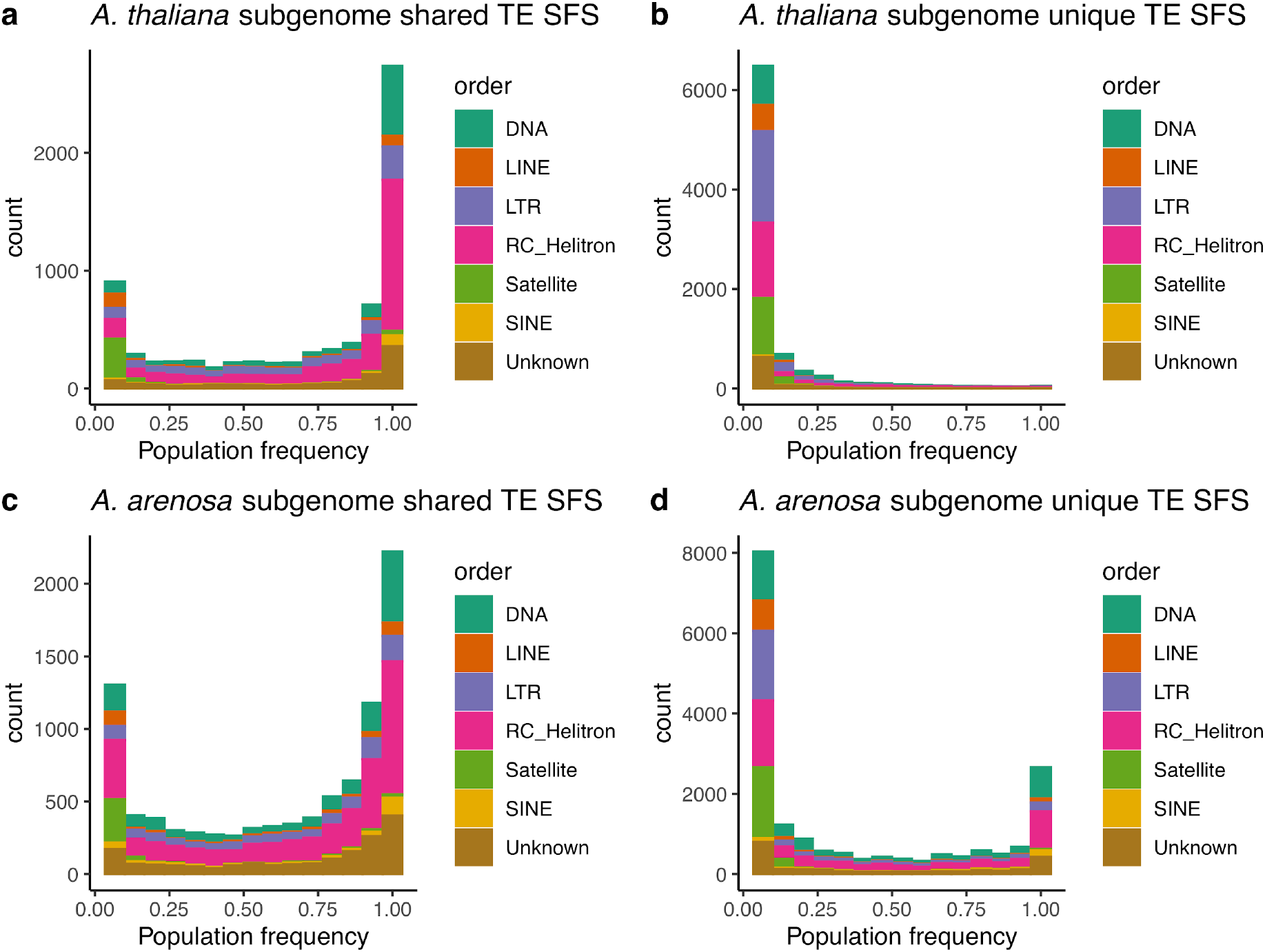
Site frequency spectrum (SFS) of shared TEs and unique TEs in *A. suecica* broken down by TE family. Shared TE SFS for the **a** *A. thaliana* and **b** *A. arenosa* subgenome. Private TE SFS for the **c** *A. thaliana* and **d** *A. arenosa* subgenome.

**Supplementary Figure 9.**
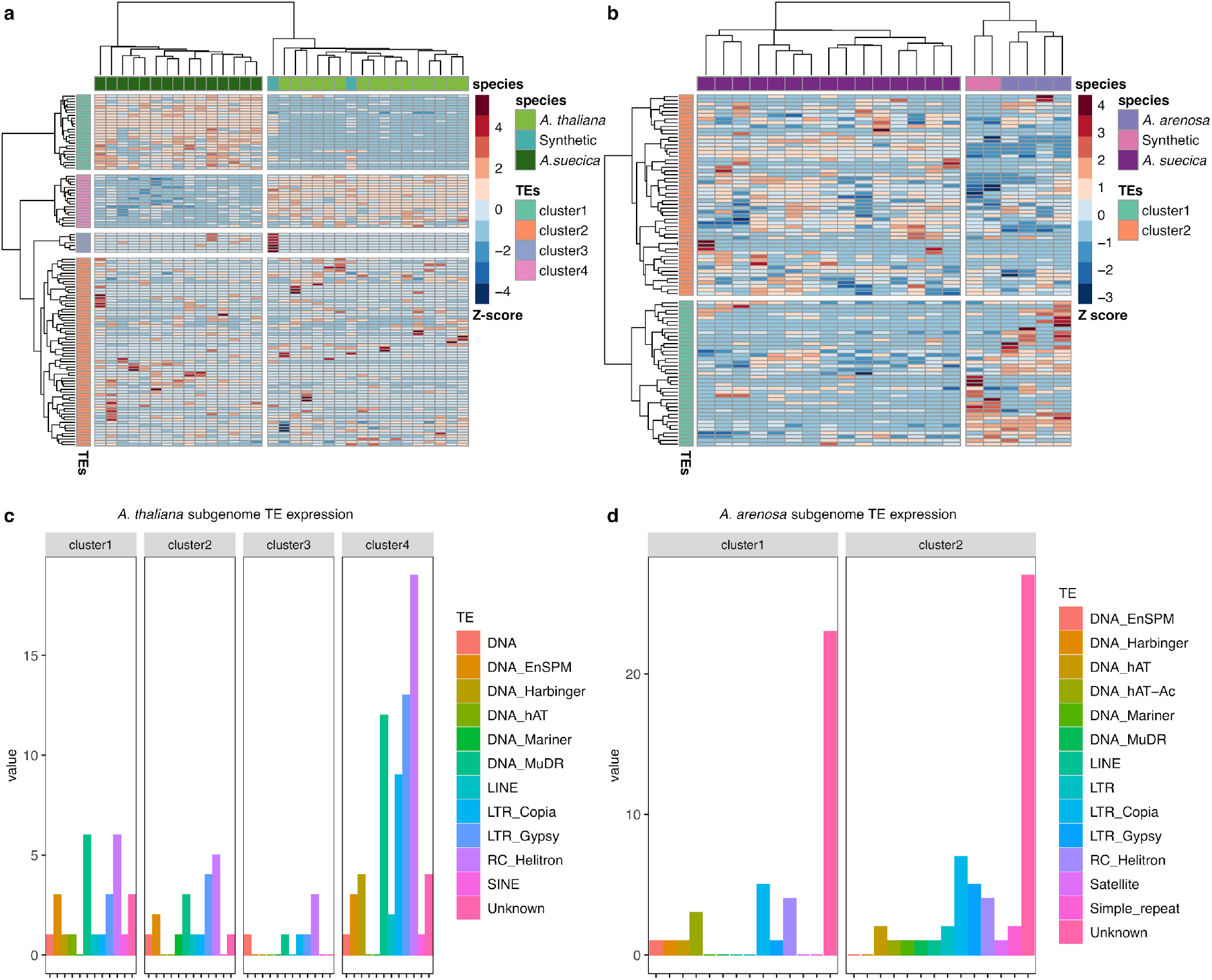
Analysis of TE expression in *A. suecica*. Patterns of TE expression in natural and synthetic *A. suecica* show that allopolyploidy is not accompanied by an overall up-regulation in TE expression as predicted by the “genome shock” hypothesis. **a** Heatmap of TE expression for the *A. thaliana* subgenome of *A. suecica* (dark green) synthetic *A. suecica* (cyan) and *A. thaliana* (light green). **b** Heatmap of TE expression for the *A. arenosa* subgenome of *A. suecica* (dark purple) synthetic *A. suecica* (pink) and *A. arenosa* (light purple). **c** and **d** the breakdown of TE families expressed in each cluster, with helitrons being the most abundant class on the *A. thaliana* subgenome and TEs of an unknown family being the most abundant in the *A. arenosa* subgenome.

**Supplementary Figure 10.**
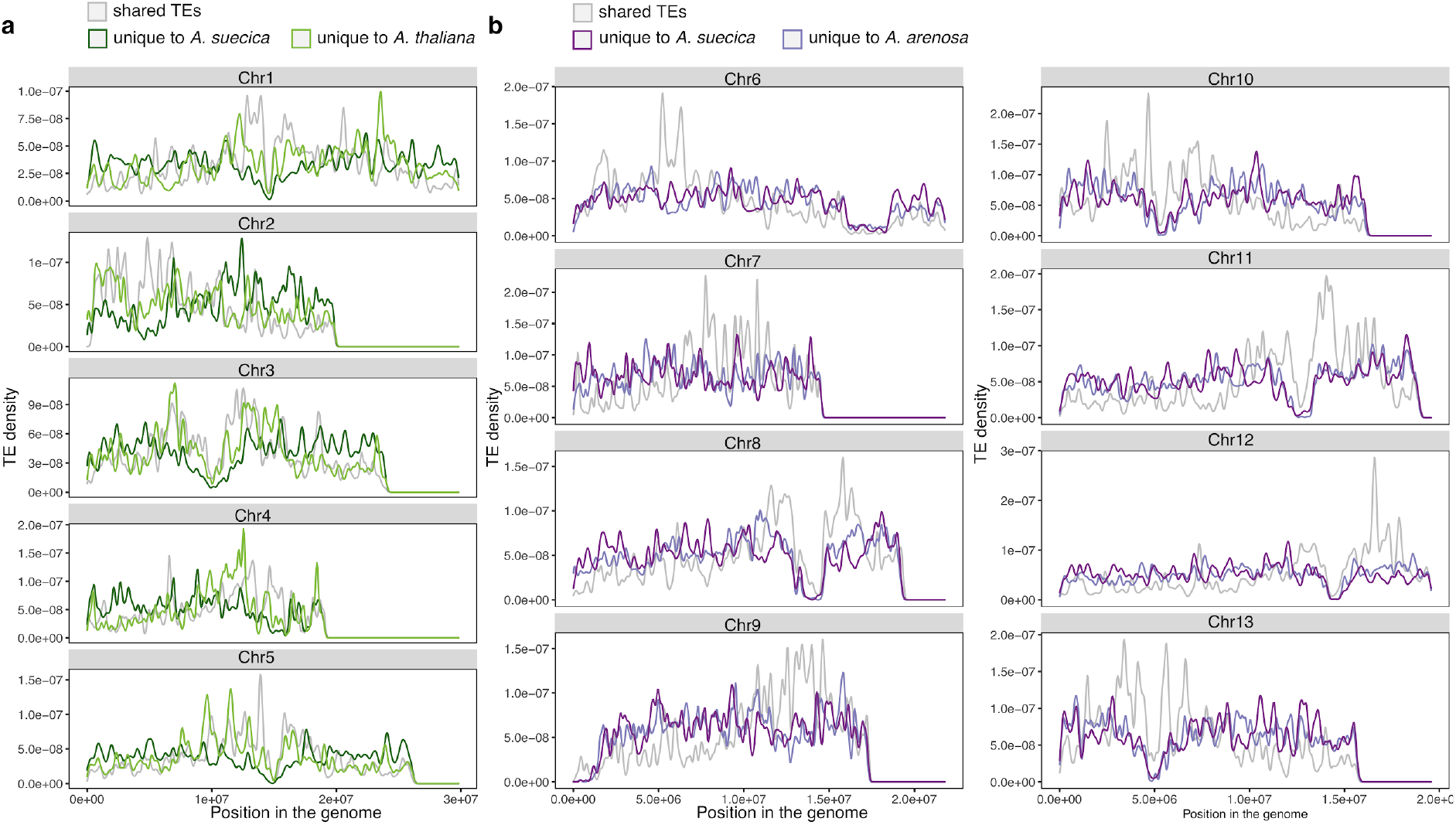
Genomic distribution of TEs in the *A. suecica* genome. **a** Shared TEs in the population between *A. thaliana* and the *A. thaliana* subgenome of *A. suecica.* Shared TEs are likely older than private TEs and are enriched around the pericentromeric regions in the *A. thaliana* subgenome. Private TEs are enriched in the chromosomal arms for both species, where protein coding gene density is higher (Fig. 1b). **b** as in **a** but examining TEs in the population of *A. arenosa* and the *A. arenosa* part of *A. suecica*. Note the region between 5 and 10 on chromosome 2 was not included in the analysis as this region shows synteny with an unplaced contig.

**Supplementary Fig 11.**
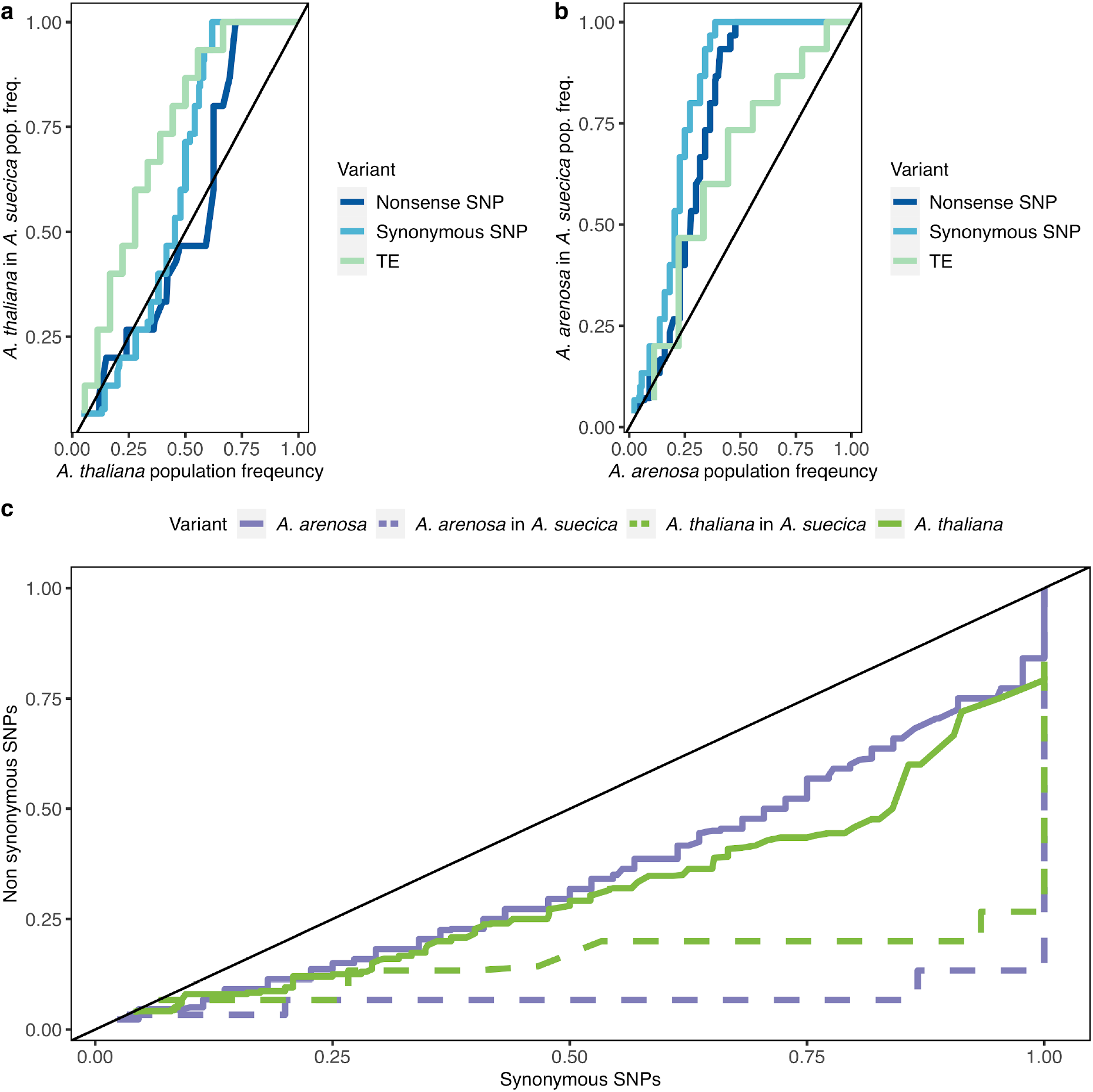
Patterns of selection in *A. suecica.* **a** Comparison of shared variation (Nonsense SNPs, synonymous SNPs, and TEs) population frequencies in the *A. thaliana* subgenome of 15 natural *A. suecica* accessions and the closest 31 *A. thaliana* accessions. **b** Comparison of shared variation (Nonsense SNPs, synonymous SNPs, and TEs) frequencies in *A. arenosa* subgenome of 15 *A. suecica* accessions and 11 Swedish *A. arenosa* lines. Although results may be affected by the sampling and potential misidentification of the ancestral populations, the current data suggests a similar pattern on both of the subgenomes for TEs and SNPs showing a bottleneck effect. **c** Plotting quantile pairs of the population frequencies of private nonsynonymous and synonymous SNPs in *A. suecica* and ancestral populations against each other, each species shows evidence of evolution under purifying selection, since population frequency quantiles of nonsynonymous SNPs are skewed to lower values than population frequency quantiles of synonymous SNPS.

**Supplementary Figure 12.**
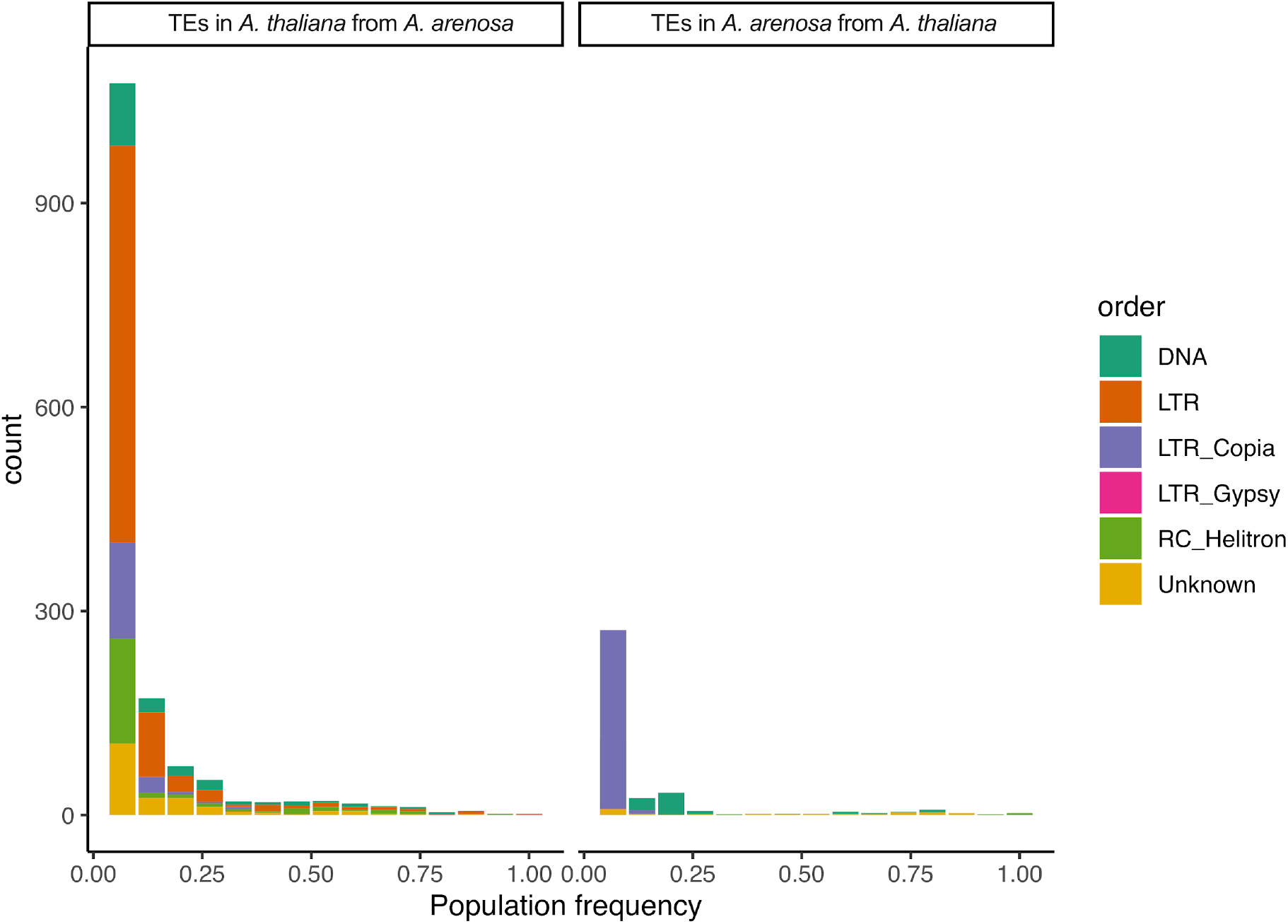
Population frequencies of presence-absence calls for TEs that have mobilized between the subgenomes in *A. suecica.* **a** TEs ancestrally from *A. arenosa* that are present in the *A. thaliana* subgenome of *A. suecica* and **b** TEs ancestrally from *A. thaliana* that are present in the *A. arenosa* subgenome of *A. suecica*.

**Supplementary Figure 13.**
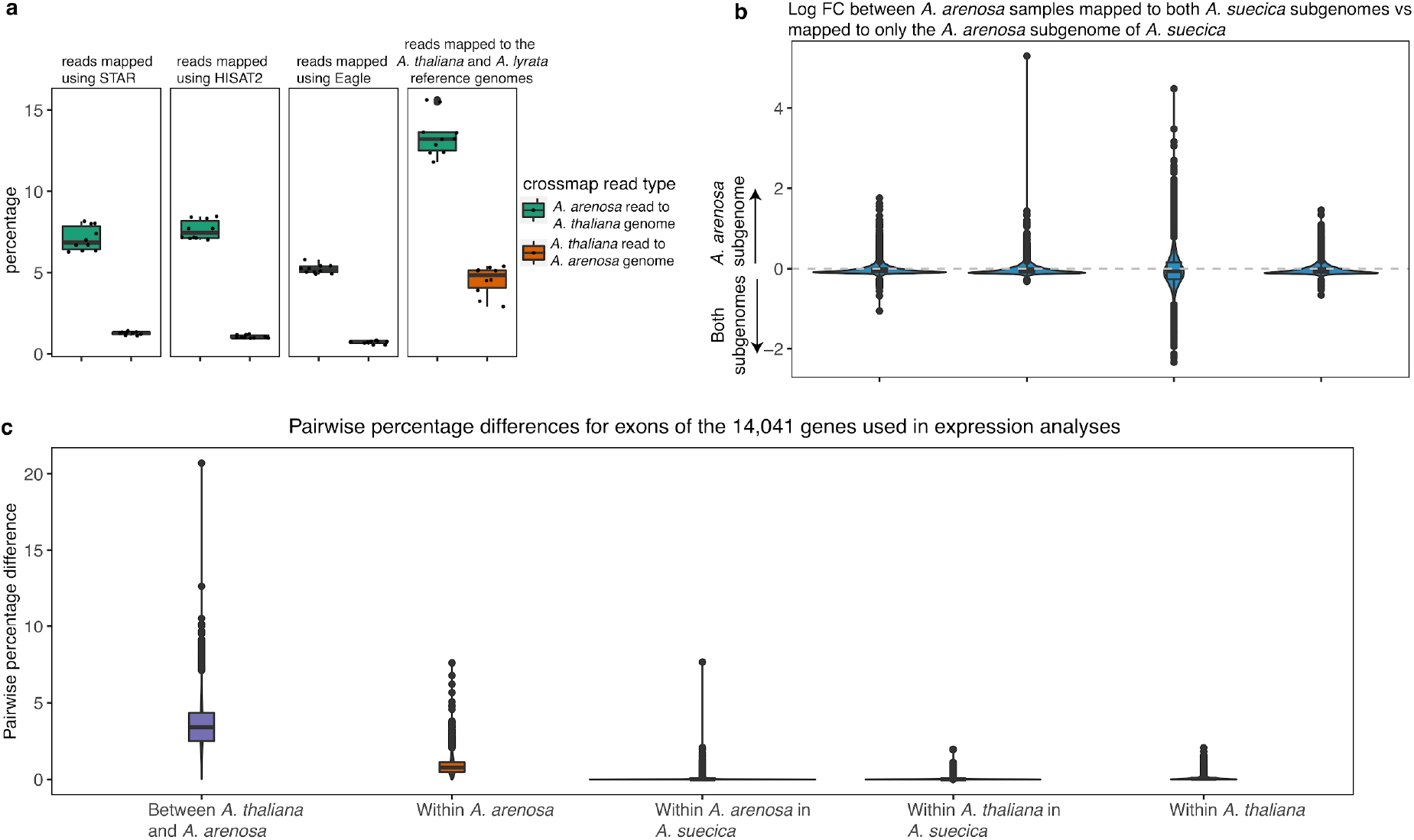
Cross-mapping in RNA-seq. **a** Boxplots of cross-mapping reads. This was examined by mixing reads in-silico between *A. thaliana* and *A. arenosa*. On average ∼6% of *A. arenosa* reads map to *A. thaliana* subgenome instead of the *A. arenosa* subgenome, and ∼1% vice versa. Mapping these reads to the combined reference genomes of *A. thaliana* and *A. lyrata* (boxplot 4 in **a**) shows that reads map more precisely to the *A. suecica* reference and that cross-mapping is not due to unreported homeologous exchange. **b** LogFC of log2 CPM read counts for *A. arenosa* (CPM of *A. arenosa* subgenome genes when reads are mapped only to *A. arenosa* subgenome of *A. suecica*/CPM of *A. arenosa* subgenome genes when reads are mapped to the full genome) show only a small effect of mapping strategy to estimate gene expression on the *A. arenosa* subgenome. **c** Pairwise percentage differences (π) for each group measured for the exons of the 14,041 genes in the expression analysis. High levels of π in *A. arenosa* overlaps with the distribution of π between *A. thaliana* and *A. arenosa.* This explains why there is more cross-mapping for *A. arenosa* than for *A. thaliana* in **a** Importantly, lower π within *A. suecica* for both subgenomes means that measurements for subgenome dominance are not biased by cross-mapping, as we expect less cross-mapping since the distribution of π overlaps less with π between *A. thaliana* and *A. arenosa*.

**Supplemental figure 14.**
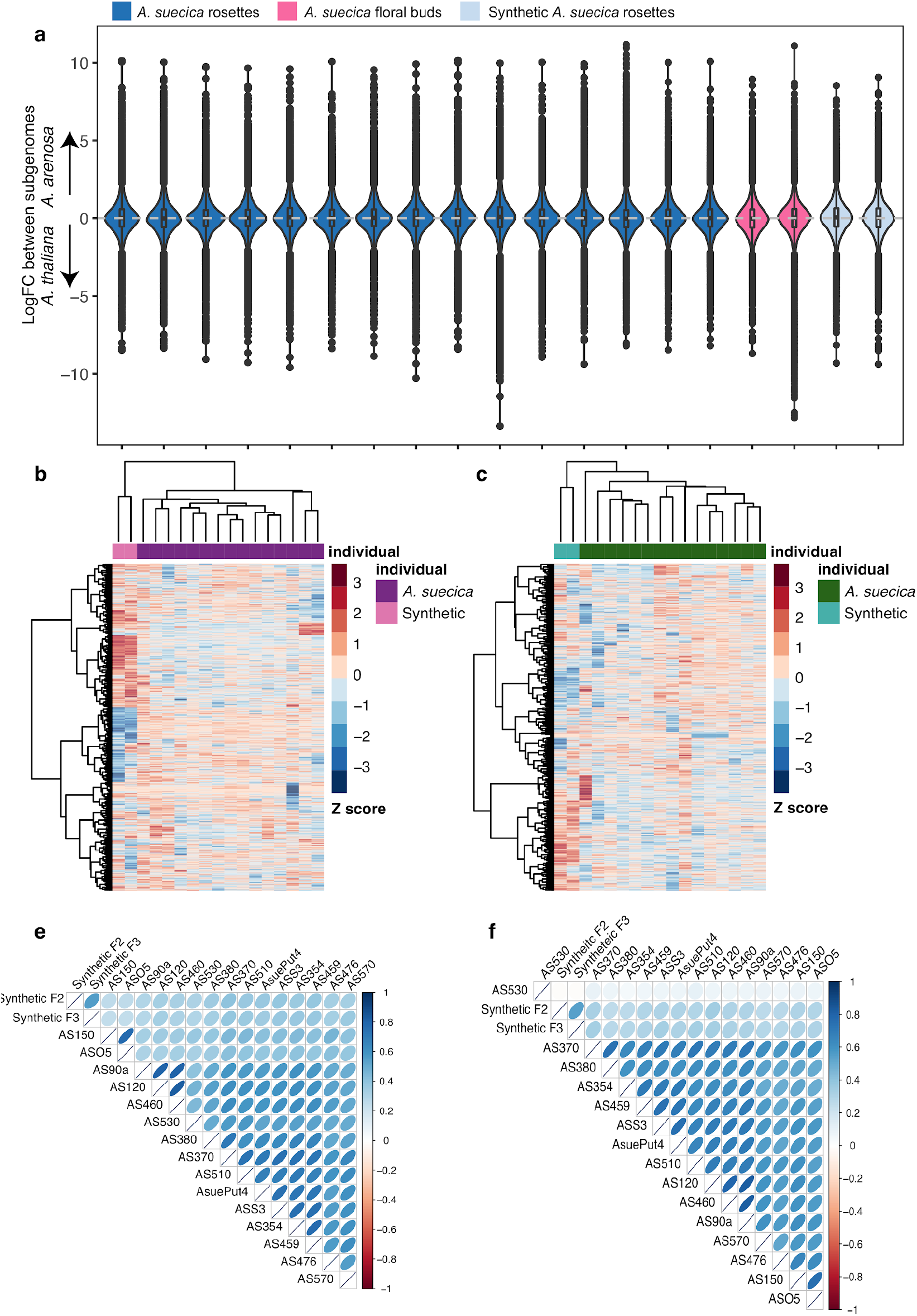
Expression differences between subgenomes in natural and synthetic *A. suecica*. **a** The distribution of expression differences across homeologous gene pairs in natural and synthetic *A. suecica*. **b** A heatmap of expression for genes in the top 5% biased toward the *A. arenosa* subgenome. The gene must be in the 5% quantile for at least 1 accession. **c** The same as in **b** but for the *A. thaliana* subgenome. Correlations of log fold change for genes in the tails of the distribution (top 5% quantile) for the *A. arenosa* subgenome **d** and the *A. thaliana* subgenome **e**

**Supplementary Figure 15.**
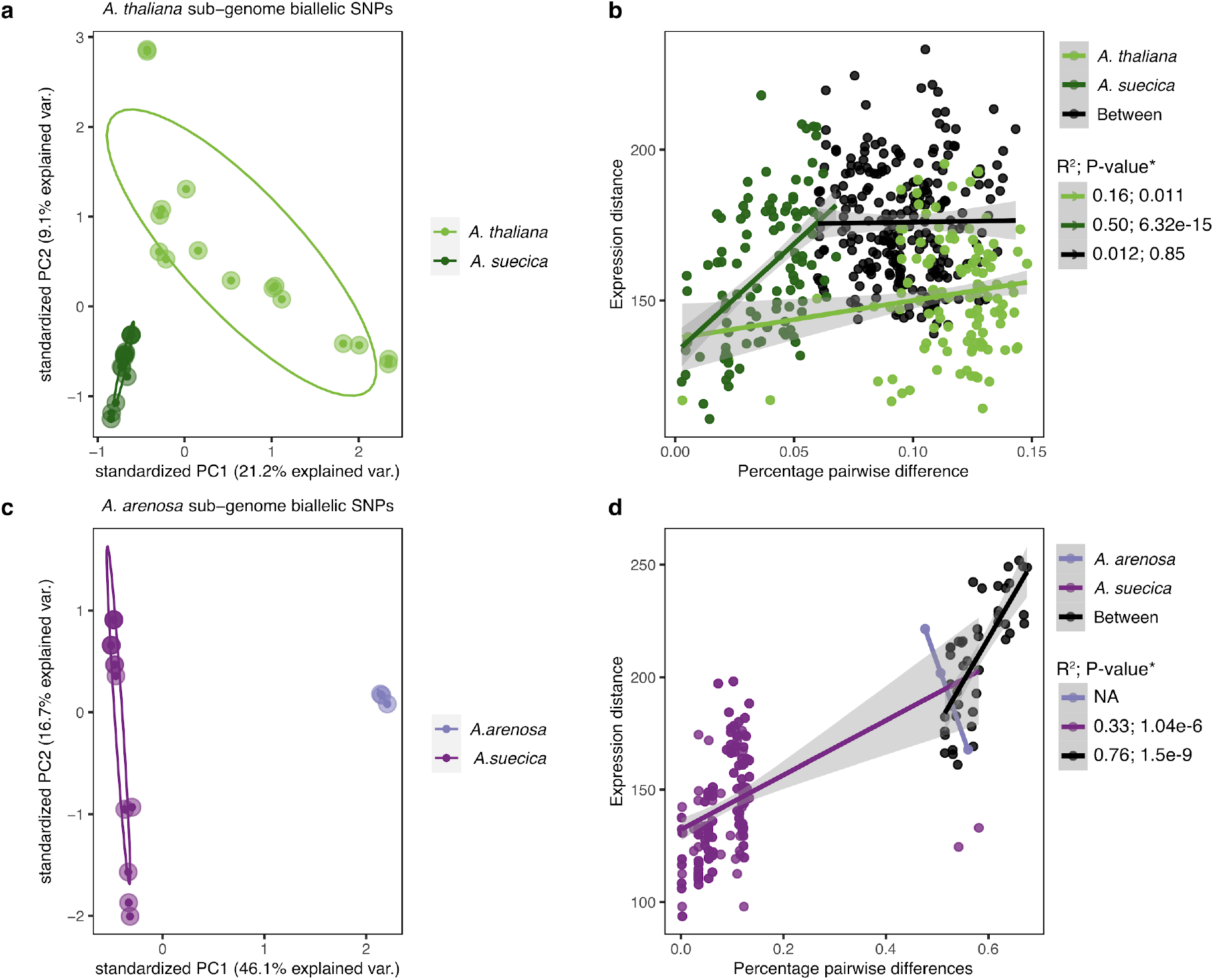
Comparison of genetic and expression distance. **a** PCA plot of biallelic SNPs in the population of *A. thaliana* and *A. suecica* for the *A. thaliana* subgenome of *A. suecica* (N=345,075 biallelic SNPs), of the analyzed 13,647 genes in gene expression in addition to 500bp up and downstream of each gene sequence **b** Correlation of *π* (pairwise genetic differences) and expression distance (i.e. euclidean distance) for 14,041 genes (*=Bootstrapped 1000 times). **c** PCA plot of biallelic SNPs in the population of *A. arenosa* (N.B. we had DNA sequencing for only 3 of the 4 accessions used in the expression analysis) and *A. suecica* for the *A. arenosa* subgenome of *A. suecica* (N= 1,761,708 biallelic SNPs), of the analyzed 14,041 genes in gene expression in addition to 500bp up and downstream of each gene sequence **d** Correlation of Pi (pairwise genetic differences for mapped genomic regions) and expression distance (i.e. euclidean distance) for 14,041 genes (*=Bootstrapped 1000 times). *A. arenosa* was too few samples to give reliable correlations and therefore is NA. Grey bars represent the 95 confidence intervals.

**Supplementary Figure 16.**
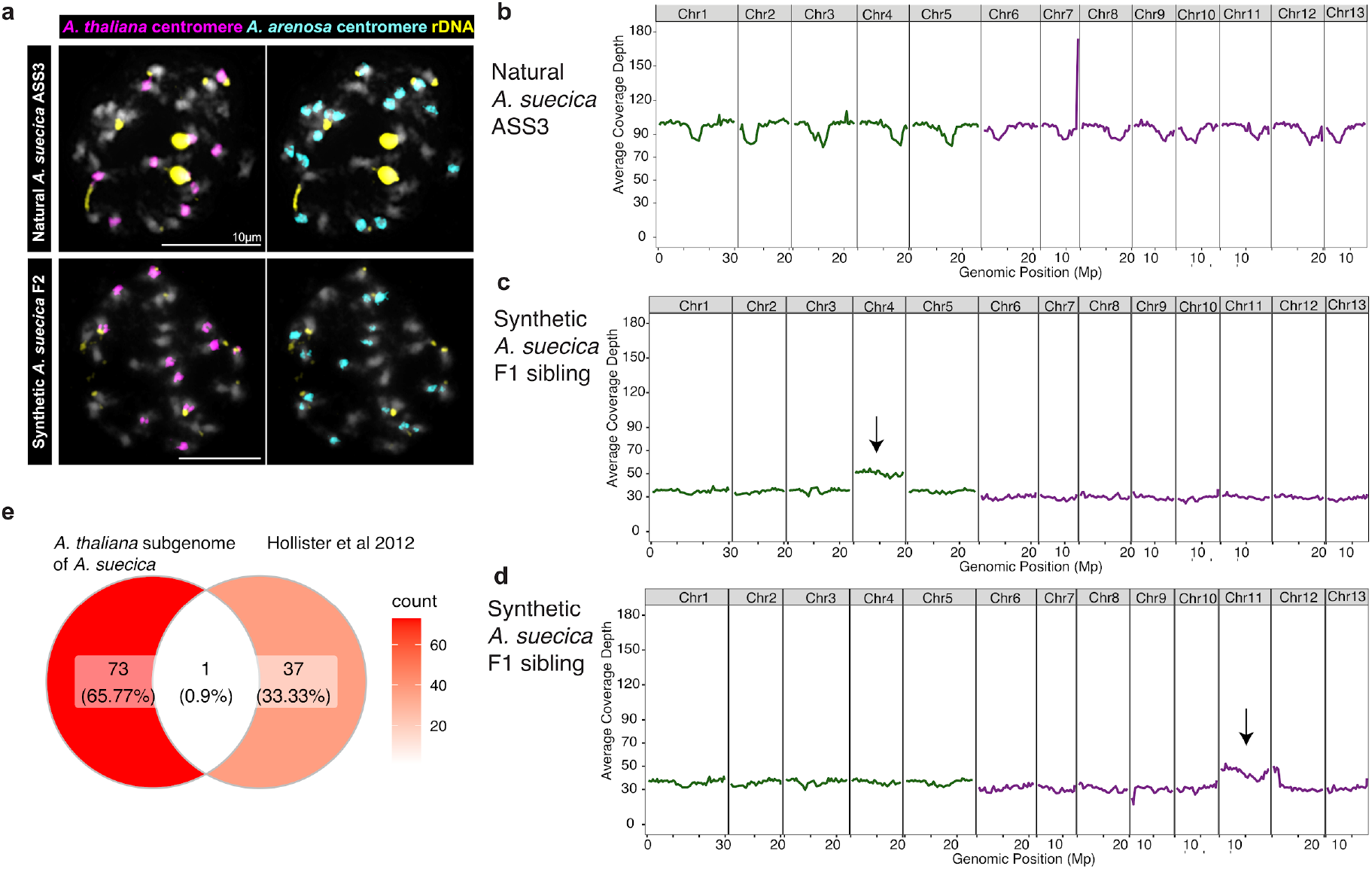
Aneuploidy is frequent in synthetic *A. suecica*. **a** Comparison of FISH analyses of the reference natural *A. suecica* “ASS3 ”and synthetic *A. suecica.* Synthetic *A. suecica* shows aneuploidy in both subgenomes in the F2 generation (gain of one chromosome on the *A. thaliana* subgenome (N=11) and loss of one chromosome on the *A. arenosa* subgenome (N=15)). Natural *A. suecica* shows a stable karyotype **b** DNA sequencing coverage in the reference natural *A. suecica* accession “ASS3” **c** and **d** DNA sequencing coverage in siblings of F1 synthetic *A. suecica* show different cases of aneuploidy (indicated with arrow) in synthetic *A. suecica*, chromosome 4 in **c** and chromosome 11 in **d e** overlap of genes involved in cell division from figure 5e and genes previously shown to play a role in the adaptation to autopolyploidy in *A. arenosa*^121^. The little overlap in genes between *A. suecica* and *A. arenosa* highlights that successful meiosis in polyploids is likely a complex trait.

**Supplementary Figure 17.**
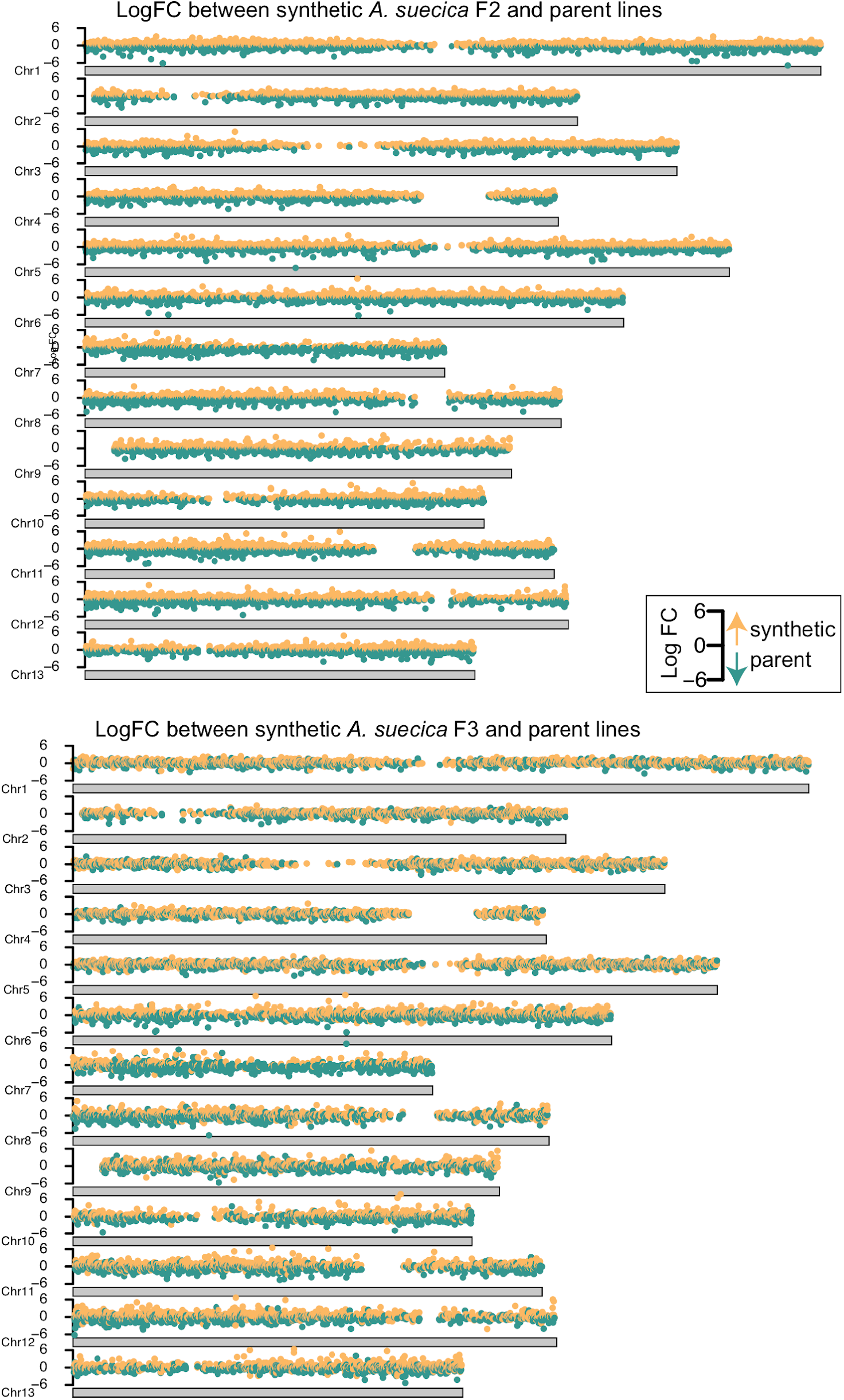
No aneuploidy in synthetic *A. suecica* lines used for RNA seq based on log fold change to parent lines. Log fold change for gene expression in **a** the 2nd and **b** the 3rd generation of synthetic *A. suecica* compared to the parent lines. No clear signal of aneuploidy (i.e. an elevated increase in expression for a full chromosome) is evident.

**Supplementary Figure 18.**
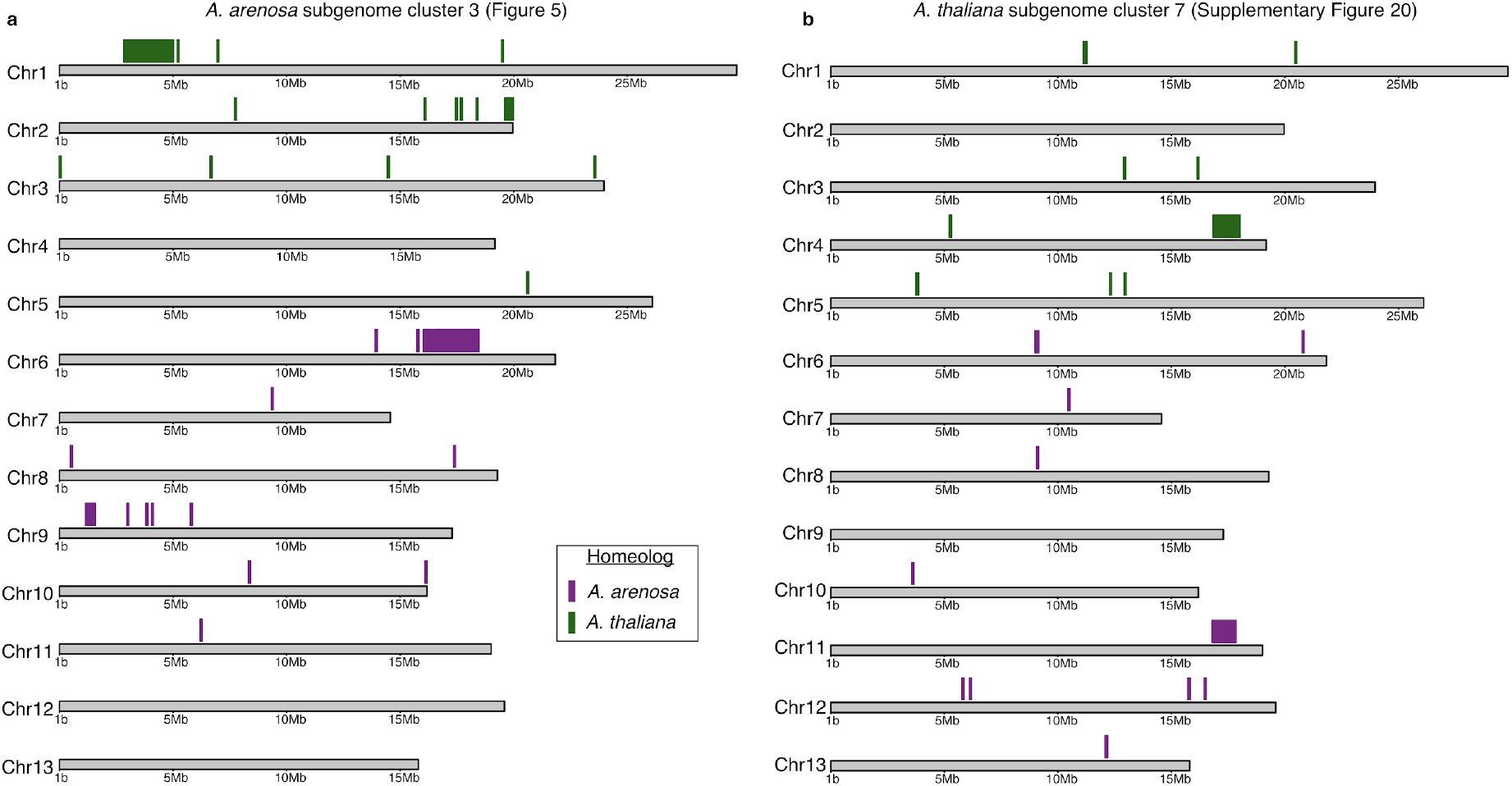
Genomic locations of genes investigated for HE signatures in *A. suecica*. **a** Genes in cluster 3 for Figure 5 in AS530 and **b** Genes in cluster 7 from Figure 18 in AS150 and ASÖ5

**Supplementary Figure 19.**
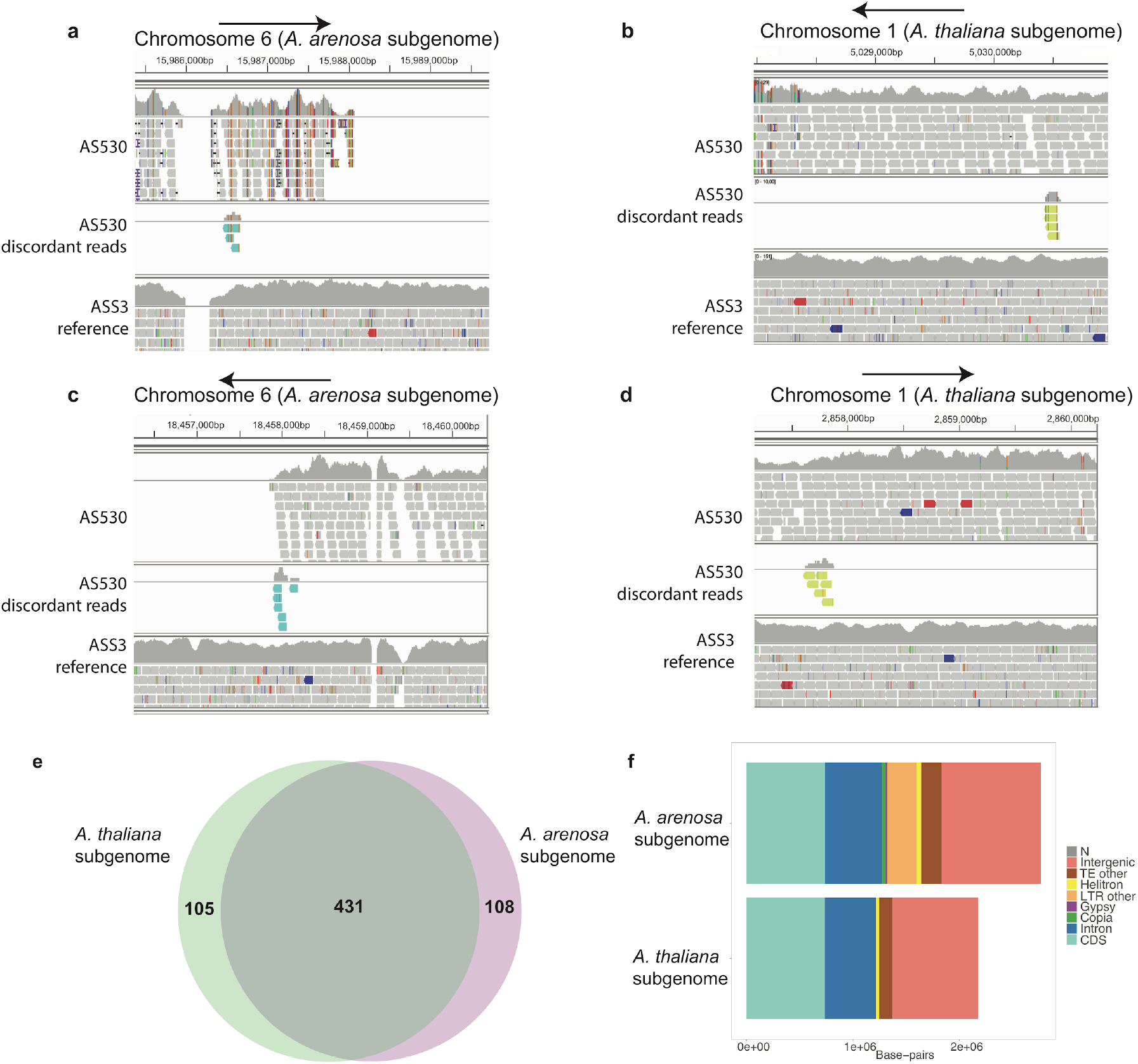
Discordant read analysis supports HE in *A. suecica*. **a** IGV screen grab of reads mapped to the beginning of the likely HE event in chromosome 6 (at ∼ 15.9Mb) before coverage depth decreases to 0 in “AS530”. Arrows point to the direction of the break along the chromosome. Discordant read pairs (cyan) map between the *A. arenosa* subgenome on chromosome 6 and the read pair (green) maps to the homeologous chromosome 1 on the *A. thaliana* subgenome (at ∼5Mb) in **b**. The end of the likely HE event in chromosome 6 (at ∼18.4Mb). Discordant reads (cyan) map between the *A. arenosa* subgenome in **c** and the read pair (green) maps to chromosome 1 (at ∼2.8Mb) on the *A. thaliana* subgenome in **d**. **e** Gene counts between the syntenic regions. 431 have a 1:1 relationship, 108 genes are specific to the *A. arenosa* subgenome in this region and 105 genes are specific to the *A. thaliana* subgenome. **f** Composition of the syntenic regions between the two subgenomes

**Supplementary Figure 20.**
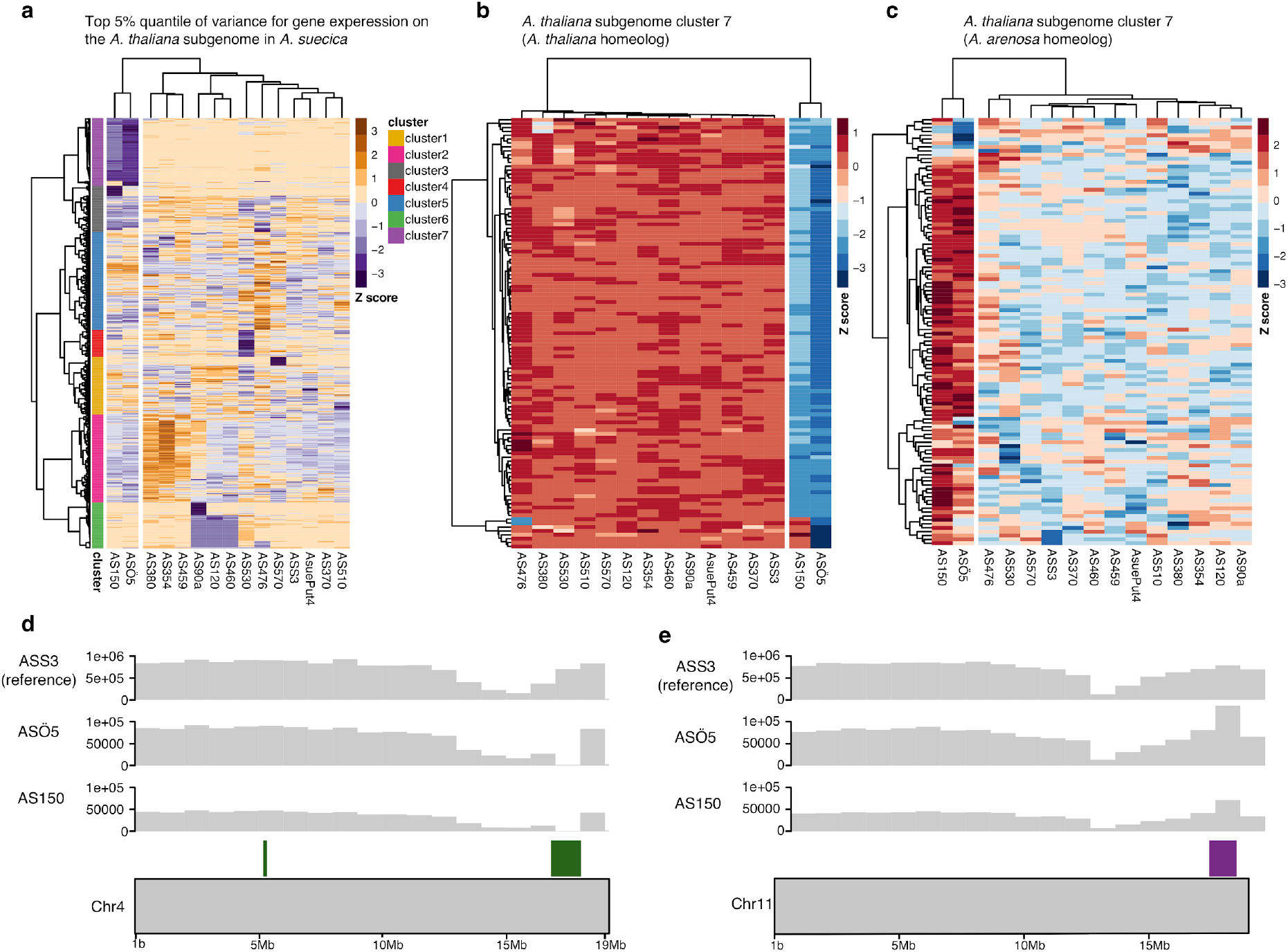
Homeologous exchange contributes to expression variance within *A. suecica* on the *A. thaliana* subgenome. **a** Taking the top 5% quantiles (N=702) for variation in gene expression for the *A. thaliana* subgenome we find a large cluster 7 (N=111) where the two outlier accessions in our PCA (“AS150” and “ASÖ5”) are expressing these genes differently to the rest of the population. **b** Homeologous genes of this cluster on the *A. thaliana* subgenome of *A. suecica* show that these genes are not expressed in these two accessions while **c** shows the opposite pattern and are higher expressed in “AS150” and “ASÖ5” compared to the rest of the population. **d** 101 of the 111 genes in cluster 7 are located on chromosome 4 in close proximity to each other on the *A. thaliana* subgenome of the *A. suecica* reference genome and appear to be deleted in “AS5Ö5” and “AS150” as they do not have DNA sequencing coverage. The *A. arenosa* subgenome homeologs (located on chromosome 11) have twice the DNA coverage, suggesting they are duplicated, in agreement with the expectations of HE event.

**Supplementary Table 1.**
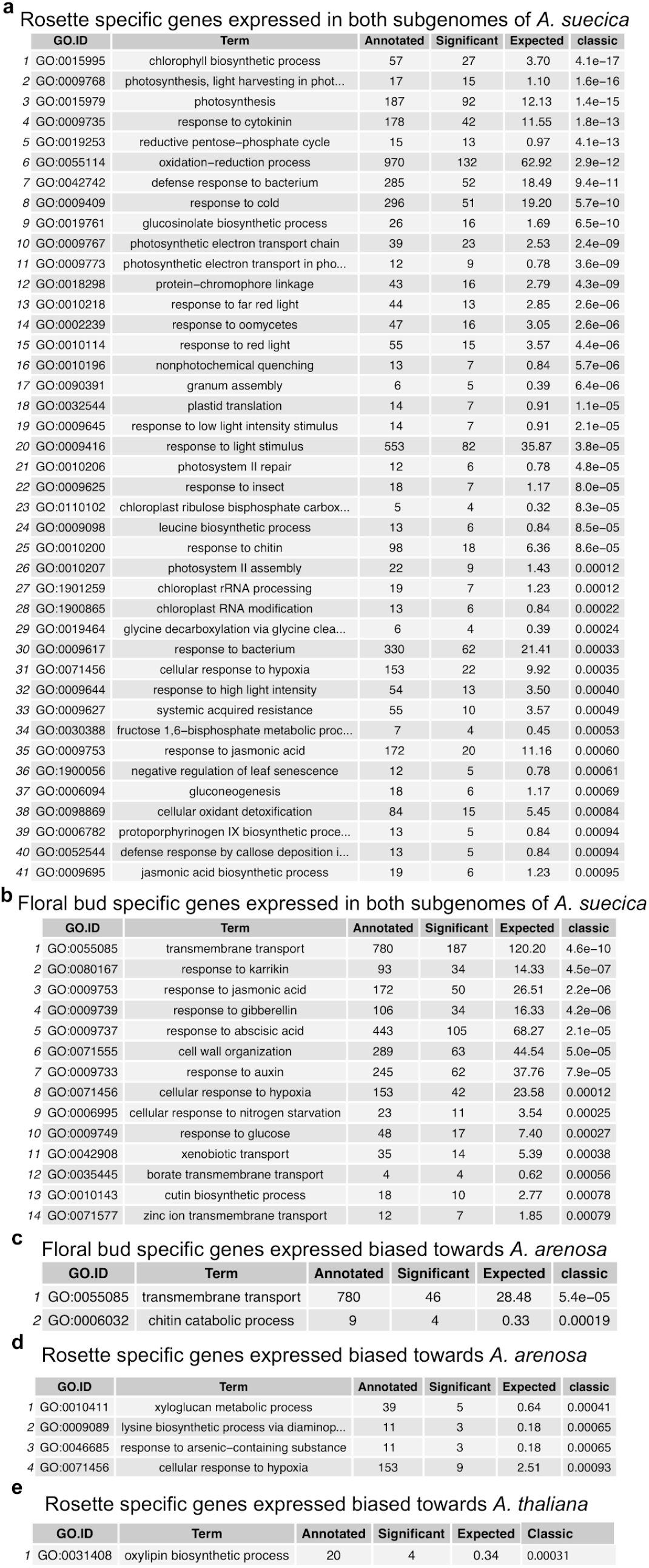
Gene ontology (GO) analysis for gene expression comparison between whole rosettes and floral buds in *A. suecica*. No significant GO was found for genes biased towards the *A. thaliana* subgenome of *A. suecica* for floral buds.

**Supplementary Table 2.**
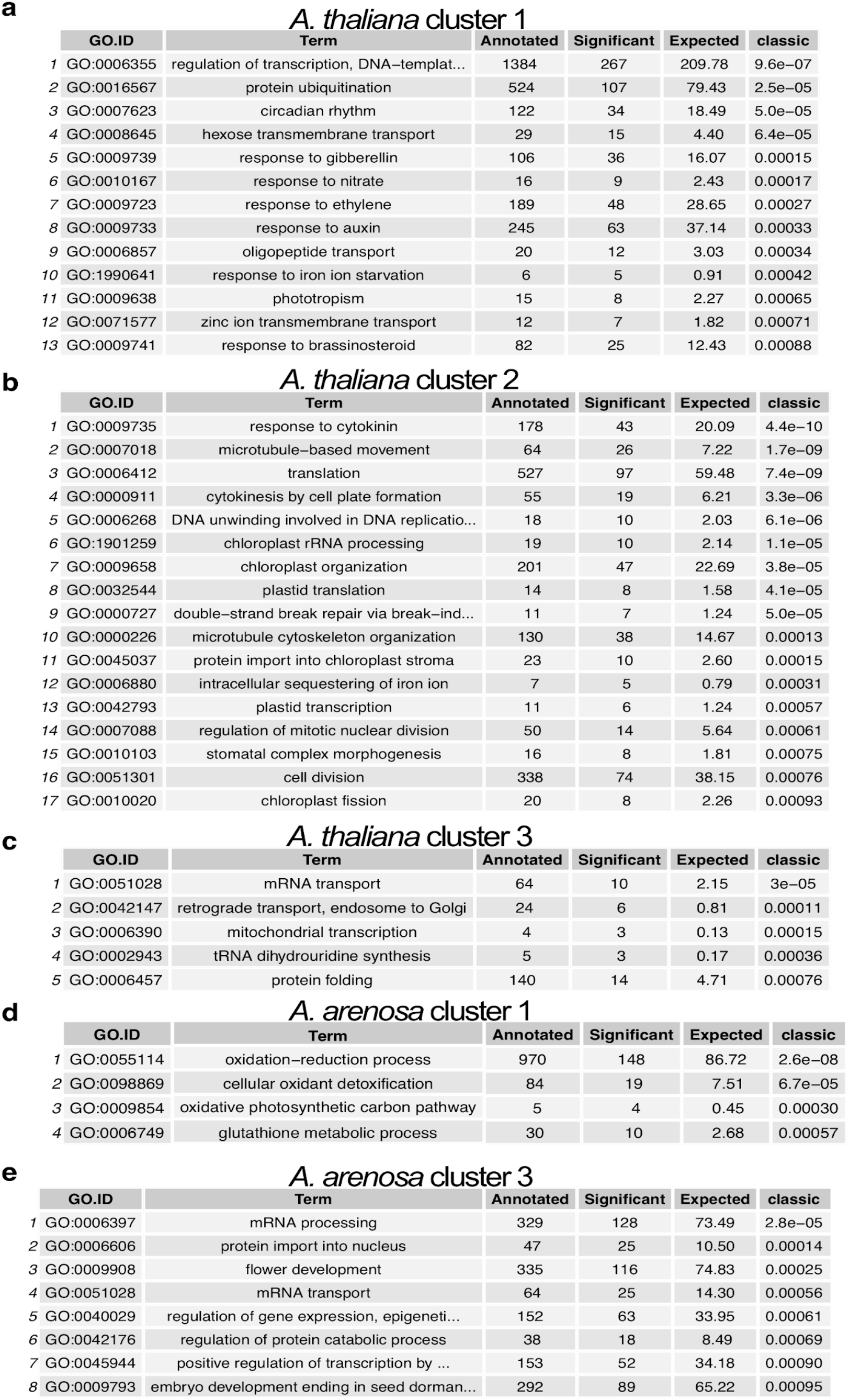
List of overrepresented gene ontologies on the Fig. 5e.

|  | GO.ID | Term | Annotated | Significant | Expected | classic |
| --- | --- | --- | --- | --- | --- | --- |
| 1 | GO:0006355 | regulation of transcription, DNA-templat... | 1384 | 267 | 209.78 | 9.6e-07 |
| 2 | GO:0016567 | protein ubiquitination | 524 | 107 | 79.43 | 2.5e-05 |
| 3 | GO:0007623 | circadian rhythm | 122 | 34 | 18.49 | 5.0e-05 |
| 4 | GO:0008645 | hexose transmembrane transport | 29 | 15 | 4.40 | 6.4e-05 |
| 5 | GO:0009739 | response to gibberellin | 106 | 36 | 16.07 | 0.00015 |
| 6 | GO:0010167 | response to nitrate | 16 | 9 | 2.43 | 0.00017 |
| 7 | GO:0009723 | response to ethylene | 189 | 48 | 28.65 | 0.00027 |
| 8 | GO:0009733 | response to auxin | 245 | 63 | 37.14 | 0.00033 |
| 9 | GO:0006857 | oligopeptide transport | 20 | 12 | 3.03 | 0.00034 |
| 10 | GO:1990641 | response to iron ion starvation | 6 | 5 | 0.91 | 0.00042 |
| 11 | GO:0009638 | phototropism | 15 | 8 | 2.27 | 0.00065 |
| 12 | GO:0071577 | zinc ion transmembrane transport | 12 | 7 | 1.82 | 0.00071 |
| 13 | GO:0009741 | response to brassinosteroid | 82 | 25 | 12.43 | 0.00088 |

b *A. thaliana* cluster 2
|  | GO.ID | Term | Annotated | Significant | Expected | classic |
| --- | --- | --- | --- | --- | --- | --- |
| 1 | GO:0009735 | response to cytokinin | 178 | 43 | 20.09 | 4.4e-10 |
| 2 | GO:0007018 | microtubule-based movement | 64 | 26 | 7.22 | 1.7e-09 |
| 3 | GO:0006412 | translation | 527 | 97 | 59.48 | 7.4e-09 |
| 4 | GO:0000911 | cytokinesis by cell plate formation | 55 | 19 | 6.21 | 3.3e-06 |
| 5 | GO:0006268 | DNA unwinding involved in DNA replicatio... | 18 | 10 | 2.03 | 6.1e-06 |
| 6 | GO:1901259 | chloroplast rRNA processing | 19 | 10 | 2.14 | 1.1e-05 |
| 7 | GO:0009658 | chloroplast organization | 201 | 47 | 22.69 | 3.8e-05 |
| 8 | GO:0032544 | plastid translation | 14 | 8 | 1.58 | 4.1e-05 |
| 9 | GO:0000727 | double-strand break repair via break-ind... | 11 | 7 | 1.24 | 5.0e-05 |
| 10 | GO:0000226 | microtubule cytoskeleton organization | 130 | 38 | 14.67 | 0.00013 |
| 11 | GO:0045037 | protein import into chloroplast stroma | 23 | 10 | 2.60 | 0.00015 |
| 12 | GO:0006880 | intracellular sequestering of iron ion | 7 | 5 | 0.79 | 0.00031 |
| 13 | GO:0042793 | plastid transcription | 11 | 6 | 1.24 | 0.00057 |
| 14 | GO:0007088 | regulation of mitotic nuclear division | 50 | 14 | 5.64 | 0.00061 |
| 15 | GO:0010103 | stomatal complex morphogenesis | 16 | 8 | 1.81 | 0.00075 |
| 16 | GO:0051301 | cell division | 338 | 74 | 38.15 | 0.00076 |
| 17 | GO:0010020 | chloroplast fission | 20 | 8 | 2.26 | 0.00093 |

c *A. thaliana* cluster 3
|  | GO.ID | Term | Annotated | Significant | Expected | classic |
| --- | --- | --- | --- | --- | --- | --- |
| 1 | GO:0051028 | mRNA transport | 64 | 10 | 2.15 | 3e-05 |
| 2 | GO:0042147 | retrograde transport, endosome to Golgi | 24 | 6 | 0.81 | 0.00011 |
| 3 | GO:0006390 | mitochondrial transcription | 4 | 3 | 0.13 | 0.00015 |
| 4 | GO:0002943 | tRNA dihydrouridine synthesis | 5 | 3 | 0.17 | 0.00036 |
| 5 | GO:0006457 | protein folding | 140 | 14 | 4.71 | 0.00076 |

d *A. arenosa* cluster 1
|  | GO.ID | Term | Annotated | Significant | Expected | classic |
| --- | --- | --- | --- | --- | --- | --- |
| 1 | GO:0055114 | oxidation-reduction process | 970 | 148 | 86.72 | 2.6e-08 |
| 2 | GO:0098869 | cellular oxidant detoxification | 84 | 19 | 7.51 | 6.7e-05 |
| 3 | GO:0009854 | oxidative photosynthetic carbon pathway | 5 | 4 | 0.45 | 0.00030 |
| 4 | GO:0006749 | glutathione metabolic process | 30 | 10 | 2.68 | 0.00057 |

|  | GO.ID | Term | Annotated | Significant | Expected | classic |
| --- | --- | --- | --- | --- | --- | --- |
| 1 | GO:0006397 | mRNA processing | 329 | 128 | 73.49 | 2.8e-05 |
| 2 | GO:0006606 | protein import into nucleus | 47 | 25 | 10.50 | 0.00014 |
| 3 | GO:0009908 | flower development | 335 | 116 | 74.83 | 0.00025 |
| 4 | GO:0051028 | mRNA transport | 64 | 25 | 14.30 | 0.00056 |
| 5 | GO:0040029 | regulation of gene expression, epigeneti... | 152 | 63 | 33.95 | 0.00061 |
| 6 | GO:0042176 | regulation of protein catabolic process | 38 | 18 | 8.49 | 0.00069 |
| 7 | GO:0045944 | positive regulation of transcription by ... | 153 | 52 | 34.18 | 0.00090 |
| 8 | GO:0009793 | embryo development ending in seed dorman... | 292 | 89 | 65.22 | 0.00095 |

## Materials & Methods

### PacBio sequencing of *A. suecica*

We used genomic DNA from whole rosettes of one *A. suecica* (“ASS3”) accession to generate PacBio sequencing data. DNA was extracted using a modified PacBio protocol for preparing *Arabidopsis* genomic DNA for size-selected ∼20kb SMRTbell libraries. Briefly, whole genomic DNA was extracted from 32g of 3-4 week old plants, grown at 16°C and subjected to a 2-day dark treatment. This generated 23 micrograms of purified genomic DNA with a fragment length of >40Kb for *A. suecica*. We assessed DNA quality with a Qubit fluorometer and a Nanodrop analysis, and ran the DNA on a gel to visualize fragmentation. Genomic libraries and single-molecule real-time (SMRT) sequence data were generated at the Functional Genomics Center Zurich (FGCZ), in Switzerland. The Pacbio RSII instrument was used with P6/C4 chemistry and an average movie length of 6 hours. A total of 12 SMRT cells were processed generating 16.3Gb of DNA bases with an N50 read length of 20 Kbp and median read length of 14 Kbp. Using the same genomic library, an additional 3.3 Gbp of data was generated by a Pacbio Sequel instrument at the Vienna Biocenter Core Facilities (VBCF), in Austria, with a median read length of 10Kbp.

### *A. suecica* genome assembly

To generate the *A. suecica* assembly we first used FALCON^127^ (version 0.3.0) with a length cutoff for seed reads set to 1 Kb in size. The assembly produced 828 contigs with an N50 of 5.81 Mb and a total assembly size of 271 Mb. Additionally, we generated a Canu^128^ (v.1.3.0) assembly using default settings, which resulted in 260 contigs with an N50 of 6.65 Mb and a total assembly size of 267 Mb. Then we merged the two assemblies using the software quickmerge^129^. The resulting merged assembly consisted of 929 contigs with an N50 of 9.02 Mb and a total draft assembly size of 276 Mb. We polished the assembly using Arrow^130^ (smrtlink release 5.0.0.6792) and Pilon (version 1.22). For Pilon^131^, 100bp (with PCR duplicates removed), and a second PCR-free 250bp, Illumina paired end reads were used that had been generated from the reference *A. suecica* accession “ASS3”.

### Pacbio sequencing of *A. arenosa*

A natural Swedish autotetraploid *A. arenosa* accession “Aa4” was inbred in a lab for two generations in order to reduce heterozygosity. We extracted whole genomic DNA from 64g of three week old plants in the same way as described for *A. suecica* (above), generating 50 μg of purified genomic DNA with a fragment sizes longer than 40 Kb in length. The *A. arenosa* genomic libraries and SMRT sequence data were generated at the Vienna Biocenter Core Facilities (VBCF), in Austria. A Pacbio Sequel instrument was used to generate a total of 22 Gbp of data from five SMRT cells, with an N50 of 13 Kbp and median read length 10 Kbp. In addition, two runs of Oxford Nanopore sequencing were carried out at the VBCF producing 750 Mbp in 180,000 reads (median 5 Kbp and 2.6 Kbp; N50 8.7 and 6.7 Kbp, respectively).

### Assembly of autotetraploid *A. arenosa*

We assembled a draft contig assembly for the autotetraploid *A. arenosa* accession “Aa4” using FALCON (version 0.3.0) as for *A. suecica*. The assembly produced 3,629 contigs with an N50 of 331 Kb, maximum contig size of 2.5 Mb and a total assembly size of 461 Mb. The assembly size is greater than the calculated haploid size of 330 Mb using FACs (see Supplementary Figure 2) probably because of the high levels of heterozygosity in *A. arenosa*. The resulting assembly was polished as described for *A. suecica*.

### HiC tissue fixation and library preparation

To generate physical scaffolds for the *A. suecica* assembly we generated proximity-ligation HiC sequencing data. We collected approximately 0.5 gram of tissue from 3-week old seedlings of the same reference *A. suecica* accession. Freshly collected plant tissue was fixed in 1% formaldehyde. Cross-linking was stopped by the addition of 0.15 M Glycine. The fixed tissue was ground to a powder in liquid nitrogen and suspended in 10 ml of nuclei isolation buffer. Nuclei was digested by adding 50 U DpnII and the digested chromatin was blunt-ended by incubation with 25 μL of 0.4 mM biotin-14-dCTP and 40 U of Klenow enzyme, as described in. 20 U of T4 DNA ligase was then added to start proximity ligation. The extracted DNA was sheared by sonication with a Covaris S220 to produce 250-500bp fragments. This was followed by size fractionation using AMPure XP beads. Biotin was then removed from unligated ends. DNA fragments were blunt-end repaired and adaptors were ligated to the DNA products following the NEBNext Ultra II RNA Library Prep Kit for Illumina.

To analyse structural rearrangements we collected tissue for 1 other natural *A. suecica* “AS530”, 1 *A. thaliana* accession ”6978”, 1 *A. arenosa* “Aa6” and 1 synthetic *A. suecica* (F3). Each sample had two replicates. We collected tissue and prepared libraries in the same manner as described above. 125bp paired-end Illumina reads were mapped using HiCUP^132^ (version 0.6.1).

### Reference-guided scaffolding of the *A. suecica* genome with *LACHESIS*

We sequenced 207 million pairs of 125bp paired-end Illumina reads from the HiC library of the reference accession “ASS3”. We mapped reads using HiCUP (version 0.6.1) to the draft *A. suecica* contig assembly. This resulted in ∼137 million read pairs with a unique alignment. Setting an assembly threshold of >= 1 Kb in size, contigs of the draft *A. suecica* assembly were first assigned to the *A. thaliana* or *A. arenosa* subgenome. To do this, we used nucmer from the software MUMmer^133^ (version 3.23) to perform whole-genome alignments. We aligned the draft *A. suecica* assembly to the *A. thaliana* TAIR10 reference and to our *A. arenosa* draft contig assembly, simultaneously. We used the MUMer command dnadiff to produce 1-to-1 alignments. As the subgenomes are only ∼86% identical, the majority of contigs could be conclusively assigned to either subgenome by examining how similar the alignments were. Contigs that could not be assigned to a subgenome based on percentage identity were examined manually, and the length of the alignment was used to determine subgenome assignment.

Finally, we used the software LACHESIS^134^ (version 1.0.0) to scaffold our draft assembly, using the reference genomes of *A. thaliana* and *A. lyrata as* a guide to assist with scaffolding the contigs (we used *A. lyrata* here instead of our draft *A. arenosa* contig assembly, as *A. lyrata* is a chromosome-level assembly). This produced a 13-scaffold chromosome-level assembly for *A. suecica*.

### Construction of the *A. suecica* genetic map

We crossed natural *A. suecica* accession “AS150” with the reference accession “ASS3". The cross was uni-directional with “AS150” as the maternal and “ASS3” as the paternal plant. F1 plants were grown, and F2 seeds were collected, from which we grew and collected 192 F2 plants. We multiplexed the samples on 96 well plates using 75bp paired end reads and generated data of 1-2x coverage per sample. Samples were mapped to the repeat-masked scaffolds of the reference *A. suecica* genome using BWA-MEM^135^ (version 0.7.15). Samtools^136^ (version 0.1.19) was used to filter reads for proper pairs and a minimum mapping quality of 5 (-F 256 -f 3 -q 5). We called variants directly from samtools mpileup output on the sequenced F2 individuals at known biallelic sites between the two accessions used to generate the cross (a total of 590,537 SNPs). We required sites to have non-zero coverage in a minimum of 20 individuals and filtered SNPs to have frequency between 0.45-0.55 in our F2 population (as the expectation is 50:50),. We removed F2 individuals that did not have genotype calls for more than 90% of the data. This resulted in 183 individuals with genotype calls for 334,257 SNPs.

Since sequencing coverage for the F2s was low this meant we had a low probability of calling heterozygous SNPs, and a higher probability of calling a SNP as homozygous. Therefore, we applied a Hidden Markov Model implemented in R package HMM^137^ to classify SNPs as homozygous or heterozygous for each of our F2 lines. We then divided the genome into 500Kb non-overlapping windows, and classified each window as homozygous (here 0 or 1, for the reference or alternate SNP) or heterozygous (here 0.5). If the frequency of 1, 0 or 0.5 represented more than 50% of the SNPs in a given window, and exceeded missing calls (NA), the window was designated as 1, 0 or 0.5 (otherwise it was NA). This was done per chromosome and the resulting file for each chromosome and their markers were processed in the R package qtl^138^, in order to generate a genetic map. Markers genotyped in less than 100 F2s were excluded from the analysis. Linkage groups were assigned with a minimum LOD score of 8 and a maximum recombination fraction of 0.35. Each chromosome was assigned to one linkage group. We defined the final marker order by the best LOD score and the lowest number of crossover events.

Notably, the assistance of a genetic map corrected the erroneous placement of a contig at the beginning of chromosome 1 of the *A. arenosa* subgenome. The misplaced contig was relocated from chromosome 1 to the pericentromeric region of chromosome 2 of the *A. arenosa* subgenome in *A. suecica*. This error was a result of a mis-assembly of chromosome 1 in the *A. lyrata* reference, as was previously pointed out ^77^. Also of note, chromosome 2 of the *A. thaliana* subgenome of *A. suecica* was previously shown to be largely devoid of intraspecific variation, thus we had sparse marker information for this chromosome in the genetic map. Therefore, this chromosome-scale scaffold was largely assembled by the manual inspection of 3D-proximity information based on our HiC sequencing and reviewing contig order using the software Juicebox^139^.

### Gene prediction and annotation of the *A. suecica* genome

We combined *de novo* and evidence-based approaches to predict protein coding genes. For *de novo* prediction, we trained AUGUSTUS^140^ on the set of conserved single copy genes using BUSCO^141^ separately on *A. thaliana* and *A. arenosa* subgenomes of *A. suecica*. The evidence-based approach included both homology to the protein sequences of the ancestral species and the transcriptome of *A. suecica*. We aligned the peptide sequences from TAIR10 *A. thaliana* assembly to the *A. thaliana* subgenome of *A. suecica*, while the peptides from *A. lyrata* from the second version of *A. lyrata* annotation^142^ (Alyrata_384_v2.1) were aligned to the *A. arenosa* subgenome of *A. suecica* using GenomeThreader^143^ (1.7.0). We mapped the RNAseq reads from the reference accession of *A. suecica* (ASS3) from the rosettes and flower buds tissues (see above) to the reference genome using tophat^144^ and generated intron hints from the split reads using bam2hints extension of AUGUSTUS. We split the alignment into *A. thaliana* and *A. arenosa* subgenomes and assembled the transcriptome of *A. suecica* for each subgenome separately in the genome-guided mode with Trinity^145^ (2.6.6). Separately for each of the subgenomes, we filtered the assembled transcripts using tpm cutoff set to 1, collapsed similar transcripts using CD-HIT^146, 147^ with sequence identity set to 90 percent, and chose the longest open reading frame from the six-frame translation. We then aligned the proteins from *A. thaliana* and *A. arenosa* parts of *A. suecica* to the corresponding subgenomes using GenomeThreader (1.7.0). We ran AUGUSTUS using retrained parameters from BUSCO and merged hints from all three sources, these being: (1) intron hints from *A. suecica* RNAseq, (2) homology hints from ancestral proteins and (3) hints from *A. suecica* proteins.

RepeatModeler^148^ (version 1.0.11) was used in order to build a *de novo* TE consensus library for *A. suecica* and identify repetitive elements based on the genome sequence. Genome locations for the identified TE repeats were determined by using RepeatMasker^149^ (version 4.0.7) and filtered for full length matches using a code described in Bailly-Bechet et. al^150^. Helitrons are the most abundant TE family in both subgenomes (Supplementary Fig. 7).

### Synthetic *A. suecica* lines

To generate synthetic *A. suecica* we crossed a natural tetraploid *A. thaliana* accession (6978 aka “Wa-1”) to a natural Swedish autotetraploid *A. arenosa* (“Aa4”) accession. Similar to the natural *A. suecica*, *A. thaliana* was the maternal and *A. arenosa* was the paternal plant in this cross. Crosses in the opposite direction were unsuccessful. We managed to obtain very few F1 hybrid plants, which after one round of selfing set higher levels of seed formation. The resulting synthetic line was able to self-fertilize. F2 seeds were descended from a common F1 and were similar to natural *A. suecica* in appearance. We further continued the synthetic line to F3 (selfed 3rd generation).

### Synteny analysis

We performed all-against-all BLASTP search using CDS sequences for the reference *A. suecica* genome and the ancestral genomes, *A. thaliana* and *A. lyrata* (here the closest substitute reference genome for *A. arenosa*, with annotation). We used the SynMap tool^151^ from the online CoGe portal^152^. We examined synteny using the default parameters for DAGChainer (maximum distance between two matches = 20 genes; minimum number of aligned pairs = 5 genes).

### Estimating copy number of rDNA repeats using short DNA reads

To measure copy number of 45S rRNA repeats in our populations of different species, we aligned short DNA reads to a single reference 45S consensus sequence of *A. thaliana*^153^. An *A. arenosa* 45S rRNA consensus sequence was constructed by finding the best hit using BLAST in our draft *A. arenosa* contig assembly. This hit matched position 1571-8232 bp of the *A. thaliana* consensus sequence, was 6,647 bp in length and is 97% identical to the *A. thaliana* 45s rRNA consensus sequence. The aligned regions of these two 45S rRNA consensus sequences, determined by BLAST, were used in copy number estimates, to ensure that the size of the sequences were equal. The relative increase in sequence coverage of these loci, when compared to the mean coverage for the reference genome, was used to estimate copy number.

### Plant material for RNA sequencing

Transcriptomic data generated in this study included 15 accessions of *A. suecica*, 16 accessions of *A. thaliana*, 4 accessions of *A. arenosa* and 2 generations of an artificial *A. suecica* line (the 2nd and 3rd selfed-generation). The sibling of a paternal *A. arenosa* parent (Aa4) and the maternal tetraploid *A. thaliana* parent (6978 aka “Wa-1”) of our artificial *A. suecica* line were included as part of our samples (Supplementary Data 1). Each accession was replicated 3 times. Seeds were stratified in the dark for 4 days at 4°C in 1 ml of sterilised water. Seeds were then transferred to pots in a controlled growth chamber at 21°C. Humidity was kept constant at 60%. Pots were thinned to 2-3 seedlings after 1 week. Pots were re-randomized each week in their trays. Whole rosettes were collected when plants reached the 7-9 true-leaf stage of development. Samples were collected between 14:00-17:00h and flash-frozen in liquid nitrogen.

### RNA extraction and library preparation

For each accession, 2-3 whole rosettes in each pot were pooled and total RNA was extracted using the ZR Plant RNA MiniPrepTM kit. We treated the samples with DNAse, and performed purification of mRNA and polyA selection using the AMPure XP magnetic beads and the Poly(A) RNA Selection Kit from Lexogen. RNA quality and degradation were assessed using the RNA Fragment Analyzer (DNF-471 stranded sensitivity RNA analysis kit, 15nt). Concentration of RNA per sample was measured using the Qubit fluorometer. Library preparation was carried out following the NEBNext Ultra II RNA Library Prep Kit for Illumina. Barcoded adaptors were ligated using NEBNext Multiplex Oligos for Illumina (Index Primers Set 1 and 2). The libraries were PCR amplified for 7 cycles. 125bp paired-end sequencing was carried out at the VBCF on Illumina (HiSeq 2500) using multiplexing.

### RNA-seq mapping and gene expression analysis

We mapped 125bp paired-end reads to the *de novo* assembled *A. suecica* reference using STAR^154^ (version 2.7), we filtered for primary and uniquely aligned reads using the parameters --outfilterMultimapNmax 1 --outSamprimaryFlag OneBestScore. We quantified reads mapped to genes using --quantMode GeneCounts.

In order to reduce signals that are the result of cross mapping between the subgenomes of *A. suecica* we used *A. thaliana* and *A. arenosa* as a control. For each gene in the *A. thaliana* subgenome we compared log fold change of gene counts in our *A. thaliana* population to those in our *A. arenosa* population. We filtered for genes with a log2(*A. thaliana*/*A. arenosa*) below 0. We applied the same filters for genes on the *A. arenosa* subgenome, here a log2(*A. arenosa*/*A. thaliana*) below 0. This reduced the number of genes analyzed from 22,383 to 21,737 on the *A. thaliana* subgenome, and 23,353 to 23,221 on the *A. arenosa* subgenome Expression analysis was then further restricted to 1:1 unique homeologous gene pairs between the subgenomes of *A. suecica* (17,881 gene pairs). Gene counts were normalized for gene size by calculating Transcripts Per Million (TPM). The effective library sizes were calculated by computing a scaling factor based on the trimmed mean of M-values (TMM) in edgeR^155^, separately for each subgenome. Lowly expressed genes were removed from the analysis by keeping genes that were expressed in at least 3 individuals of *A. thaliana* and *A. suecica*, at least 1 individual of *A. arenosa* and at least 1 individual of synthetic *A. suecica*. 14,041 homeologous gene pairs satisfied our expression criteria. Since *A. suecica* is expressing both subgenomes, in order to correctly normalize the effective library size in *A. suecica* accessions, the effective library size was calculated as a mean of TPM counts for both subgenomes. The effective library size of *A. thaliana* accessions was calculated for TPM counts using the *A. thaliana* subgenome of the reference genome, as genes from this subgenome will be expressed in *A. thaliana*, and the effective library size of *A. arenosa* lines using the *A. arenosa* subgenome of the reference *A. suecica* genome. Gene counts were transformed to count per million (CPM) with a prior count of 1, and were log2-transformed. We used the mean of replicates per accession for downstream analyses.

To compare homeologous genes between the subgenomes in *A. suecica* we computed a log-fold change using log2(*A. arenosa* homeolog/*A. thaliana* homeolog). For tissue-specific genes we took genes that showed a log-fold change >=2 in expression between two tissues. For comparing homologous genes between the (sub-)genomes of *A. suecica* and the ancestral species *A. thaliana* and *A. arenosa*, we performed a Wilcoxon test independently for each of the 14,041 homeologous gene-pairs. Using the normalised CPM values, we compared the relative expression level of a gene on the *A. thaliana* subgenome between our population of *A. thaliana* and *A. suecica*. We performed the same test on the *A. arenosa* subgenome comparing relative expression of a gene between our population of *A. arenosa* and *A. suecica*. We filtered for genes with an adjusted p-value below <0.05 (using FDR correction). This amounted to 4,186 and 4,571 DEGs for the *A. thaliana* and *A. arenosa* subgenomes, respectively.

Cross-mapping between subgenomes was measured by mixing RNA reads between *A. thaliana* and *A. arenosa* and mapping to the *A. suecica* genome. ∼1% of A. thaliana reads map to the A. arenosa subgenome and ∼6% of the *A. arenosa* reads map to the *A. thaliana* subgenome, regardless of mapping strategy or pipeline (see Supplementary Figure 13). This can be explained by pairwise percentage differences or π within *A. arenosa* overlapping this distribution of π between *A. thaliana* and *A. arenosa* such that some exons on the *A. thalian*a subgenome are in fact closer to a particular *A. arenosa* individual than those on the *A. arenosa* subgenome of *A. suecica*. However lower π in *A. suecica* suggest this observation will not affect estimates of subgenome dominance for *A. suecica*.

### Expression analysis of rRNA

RNA reads were mapped in a similar manner as DNA reads for the analysis of rDNA copy number (above). Expression analysis was performed in a similar manner to protein coding genes, in edgeR. We defined the exclusive expression of a particular 45S rRNA gene by taking a cut-off of 15 for log2(CPM) as this was the maximum level of cross-mapping we observed for the ancestral species (see Supplementary Fig. 6).

### Expression analysis of transposable elements

To analyse the expression of transposable elements between species, the annotated TE consensus sequences in *A. suecica* were aligned using BLAST all vs all. Highly similar TE sequences (more than 85% similar for more than 85% percent of the TE sequence length), were removed, leaving 813 TE families out of 1213. Filtered *A. suecica* TEs were aligned to annotated *A. thaliana* (TAIR10) and *A. arenosa* (the PacBio contig assembly presented in this study) TE sequences to assign each family to an ancestral species using BLAST. 208 TE families were assigned to the *A. thaliana* parent and 171 TE families were assigned to the *A. arenosa* parent.

RNA reads were mapped to TE sequences using a similar approach as for gene expression analysis using edgeR. TEs that showed expression using a cut-off of log2CPM > 2 were kept. 121 *A. thaliana* TE sequences and 93 *A. arenosa* TE sequences passed this threshold. We took the mean of replicates per accession for further downstream analyses.

### Gene ontology (GO) enrichment analysis

We used the R package TopGO^156^ to conduct gene ontology enrichment analysis. We used the “weight01” algorithm when running TopGO which accounts for the hierarchical structure of GO terms and thus implicitly corrects for multiple testing. GO annotations were based on the *A. thaliana* ortholog of *A. suecica* genes. Gene annotations for *A. thaliana* were obtained using the R package biomaRt^157^ from Ensembl ‘biomaRt::useMart(biomart = “plants_mart", dataset = “athaliana_eg_gene", host = ’plants.ensembl.org’).

### Genome sizes measurements

We measured genome size for the reference *A. suecica* accession “ASS3” and the *A. arenosa* accession used for PacBio “Aa4”, using *Solanum lycopersicum* cv. Stupicke (2C = 1.96 pg DNA) as the standard. The reference *A. lyrata* accession “MN47” and the *A. thaliana* accession “CVI” were used as additional controls. Each sample had 2 replicates.

In brief, the leaves from three week old fresh tissue were chopped using a razor blade in 500 µl of UV Precise P extraction buffer + 10 µl mercaptoethanol per ml (kit PARTEC CyStain PI Absolute P no. 05-5022) to isolate nuclei. Instead of the Partec UV Precise P staining buffer, however, 1 ml of a 5 mg DAPI solution was used, as DAPI provides DNA content histograms with high resolution. The suspension was then passed through a 30 µm filter (Partec CellTrics no. 04-0042-2316) and incubated for 15 minutes on ice before FACs.

Genome size was measured using flow cytometry and a FACS Aria III sorter with near UV 375nm laser for DAPI. Debris was excluded by selecting peaks when plotting DAPI-W against DAPI-A for 20,000 events.

The data were analyzed using the flowCore^158^ package in R. Genome size was estimated by comparing the mean G1 of the standard *Solanum lycopersicum* to that of each sample to calculate the 2C DNA content of that sample using the equation:

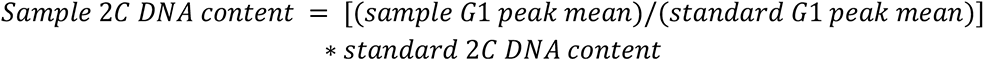

We also measured genome size for the reference *A. suecica* accession “ASS3” using the software jellyfish^159^ and findGSE^160^ using kmers (21mers). The genome size estimated was 312Mb, compared to the 305Mb estimated using FACs (see Supplementary Fig 1).

### Mapping of TE insertions

We used PopoolationTE2^100^ (version v1.10.04) to identify TE insertions. The advantage of this TE-calling software to others is that it avoids a reference bias by treating all TEs as *de-novo* insertions. Briefly, it works by using discordant read pairs to calculate the location and abundance of a TE in the genome for an accession of interest.

We mapped 100 bp Illumina DNA reads from ^20, 76, 161^, in addition to our newly generated synthetic *A. suecica* using BWA MEM^135^ (version 0.7.15) to a repeat-masked version of the *A. suecica* reference genome, concatenated with our annotated repeat sequences (see ‘Genome annotation’), as this is the data format required by PopoolationTE2. Reads were given an increased penalty of 15 for being unpaired. Reads were de-duplicated using Samtools^136^ rmdup (version 1.9). The resulting bam files were then provided to PopoolationTE2 to identify TE insertions in the genome of each of our *A. suecica*, *A. thaliana* and *A. arenosa* accessions. We used a mapping quality of 10 for the read in the discordant read pair mapping to the genome. We used the ‘separate’ mode in the ‘identify TE signatures’ step and a ‘--min-distance -200 --max-distance 500’ in the ‘pairupsignatues’ step of the pipeline. TE counts within each accession were merged if they fell within 400 bp of each other and if they mapped to the same TE sequence. All TE counts (i.e. the processed TE counts for each accession) were then combined to produce a population-wide count estimate. Population wide TE insertions were merged if they mapped to the same TE sequence and fell within 400 bp of each other. Coverage of each TE insertion in the population was also calculated for each accession. The final file was a list TE insertions present in the population and the presence or absence (or “NA” if there was no coverage to support the presence or absence of a TE insertion) in each accession analyzed (Supplementary Data 1).

### Assigning ancestry to TE sequences

In order to examine TE consensus sequences that have mobilized between the subgenomes of *A. suecica*, we first examined which of our TE consensus sequences (N=1152) have at least the potential to mobilize (i.e. have full length TE copies in the genome of *A. suecica)*. We filtered for TE consensus sequences that had TE copies in the genome of *A. suecica* that are more than 80% similar in identity for more than 80% of the consensus sequence length (N=936). Of these, 188 consensus sequences were private to the *A. thaliana* subgenome, 460 were private to the *A. arenosa* subgenome, and 288 TE consensus sequences were present in both subgenomes of *A. suecica*. To determine if TEs have jumped from the *A. thaliana* subgenome to the *A. arenosa* subgenome and vice versa we next needed to assign ancestry to these 288 TE consensus sequences. To do this we used BLAST to search for these consensus sequences in the ancestral genomes of *A. suecica*, using the TAIR10 *A. thaliana* reference and our *A. arenosa* PacBio contig assembly. Using the same 80%-80% rule we assigned 55 TEs to *A. arenosa* and 15 TEs to *A. thaliana* ancestry.

### Read mapping and SNP calling

To call biallelic SNPs we mapped reads to the *A. suecica* reference genome using the same filtering parameters described in “Mapping of TE insertions”. Biallelic SNPs were called using HaplotypeCaller from GATK^162^ (version 3.8) using default quality thresholds. SNPs were annotated using SnpEff^163^. Biallelic SNPs on the *A. thaliana* sub-genome were polarized using 38 diploid *A. lyrata* lines^76^ and biallelic SNPS on the *A. arenosa* sub-genome were polarized using 30 *A. thaliana* accessions^161^ closely related to *A. suecica*^20^.

### Chromosome preparation and FISH

Whole inflorescences of *A. arenosa*, *A. suecica* and *A. thaliana* were fixed in freshly prepared ethanol:acetic acid fixative (3:1) overnight, transferred into 70% ethanol and stored at -20°C until use. Selected inflorescences were rinsed in distilled water and citrate buffer (10 mM sodium citrate, pH 4.8), and digested by a 0.3% mix of pectolytic enzymes (cellulase, cytohelicase, pectolyase; all from Sigma-Aldrich) in citrate buffer for c. 3 hrs. Mitotic chromosome spreads were prepared from pistils as previously described^164^ by Mandáková and Lysak and suitable slides pretreated by RNase (100 µg/ml, AppliChem) and pepsin (0.1 mg/ml, Sigma-Aldrich).

For identification of *A. thaliana* and *A. arenosa* subgenomes in the allotetraploid genome of *A. suecica*, FISH probes were made from plasmids pARR20–1 or pAaCEN containing 180 bp of *A. thaliana* (pAL; Vongs et al. 1993) or ∼250 bp of *A. arenosa* (pAa; Kamm et al. 1995) pericentromeric repeats, respectively. The *A. thaliana* BAC clone T15P10 (AF167571) bearing 45S rRNA gene repeats was used for in situ localization of NORs. Individual probes were labeled with biotin-dUTP, digoxigenin-dUTP and Cy3-dUTP by nick translation, pooled, precipitated, and resuspended in 20 µl of hybridization mixture [50% formamide and 10% dextran sulfate in 2× saline sodium citrate (2× SSC)] per slide as previously described^96^.

Probes and chromosomes were denatured together on a hot plate at 80°C for 2 min and incubated in a moist chamber at 37°C overnight. Post hybridization washing was performed in 20% formamide in 2× SSC at 42°C. Fluorescent detection was as follows: biotin-dUTP was detected by avidin–Texas Red (Vector Laboratories) and amplified by goat anti-avidin–biotin (Vector Laboratories) and avidin–Texas Red; digoxigenin-dUTP was detected by mouse anti-digoxigenin (Jackson ImmunoResearch) and goat anti-mouse Alexa Fluor 488 (Molecular Probes). Chromosomes were counterstained with DAPI (4’,6-diamidino-2-phenylindole; 2 μg/ml) in Vectashield (Vector Laboratories). Fluorescent signals were analyzed and photographed using a Zeiss Axioimager epifluorescence microscope and a CoolCube camera (MetaSystems). Images were acquired separately for the four fluorochromes using appropriate excitation and emission filters (AHF Analysentechnik). The monochromatic images were pseudo colored and merged using Adobe Photoshop CS6 software (Adobe Systems).

### DAP-seq enrichment analysis for transcription factor target genes

We downloaded the target genes of transcription factors from the plant cistrome database (http://neomorph.salk.edu/dap_web/pages/index.php), which is a collection of transcription factor binding sites and their target genes, in *A. thaliana*, based on DAP-seq^165^. To test for enrichment of a gene set (for example the genes in *A. thaliana* cluster 2 on Fig. 5) for target genes of a particular transcription factor, we performed a hyper-geometric test in R. As a background we used the total 14,041 genes used in our gene expression analysis. We then performed FDR correction for multiple testing to calculate an accurate p-value of the enrichment.

### Data Availability

Genome assemblies and raw short reads can be found in the European Nucleotide Archive (ENA) (https://www.ebi.ac.uk/ena/browser/home).

The genome assembly for *A. suecica* ASS3 can be found under the BioProject number PRJEB42198, assembly accession GCA_905175345. The raw reads for the *A. suecica* genome assembly generated by Pacbio RSII can be found under ERR5037702 and those from Sequel under ERR5031296. The HiC reads used for scaffolding the *A. suecica* assembly can be found under ERR5032369.

The contig assembly for tetraploid *A. arenosa (ssp. arenosa)* can be found under the BioProject number PRJEB42276, assembly accession GCA_905175405. The raw reads for the *A. arenosa* Aa4 contig assembly generated by Sequel can be found under ERR5031542 and the reads generated by Nanopore under ERR5031541. HiC reads for the *A. arenosa* assembly can be found under ERR5032370.

HiC sequencing data for the ancestral species, the outlier accession AS530 and synthetic *A. suecica* can be found under the BioProject PRJEB42290.

DNA resequencing of synthetic *A. suecica* and parents generated in this study can be found under the BioProject PRJEB42291.

The RNA-seq reads are under the BioProject number PRJEB42277.

TE presence/absence calls for *A. suecica* and the ancestral species can be found in Supplementary Data 1.

A list of DEGs, orthologs, enriched DAP-seq transcription factors, CyMIRA gene overlaps and RNA-seq mapping statistics can be found in Supplementary Data 2.

Log fold change and CPM (counts per million) for genes on the *A. thaliana* and *A. arenosa* subgenome can be found in Supplementary Data 3.

The gene annotation (gff3 file) of the *A. suecica* genome can be found in Supplementary Data 4.

TE consensus sequences and a hierarchy file of TE order for *A. suecica* can be found in Supplementary Data 5.

## Acknowledgments

This work was supported, in part, by DFG SPP 1529 to M.N. and Detlef Weigel. T.M. and M.A.L. were supported by the Czech Science Foundation (grant no. 19-03442S) and the CEITEC 2020 project (grant no. LQ1601). P.Y.N. acknowledges postdoctoral fellowship of the Research Foundation–Flanders (12S9618N). We thank the Next Generation Sequencing Unit of the Vienna Biocenter Core Facilities (VBCF) for assistance. We thank Svante Holm and Torbjörn Säll for material collections and helpful discussions throughout. We also thank Yves Van de Peer for providing useful feedback on the manuscript, and Joel Sharbrough for pointing us to the CyMIRA database

